# Phosphorylation enables progressive microtubule-associated protein proteolysis and functionalisation during neural development

**DOI:** 10.1101/2025.09.19.677315

**Authors:** Zedeng Yang, Magda Liczmanska, Elizabeth K.J. Hogg, Iona Wallace, Houjiang Zhou, Robert Gourlay, Iolo Squires, Ziyi Li, Ying Hao, Fiona Brown, Rachel Toth, Thomas Macartney, C. James Hastie, Yu Andy Qi, Marek Gierlinski, Francisco Bustos, Greg M. Findlay

## Abstract

Neurological diseases are frequently characterised by dysregulation of the microtubule cytoskeleton, which is critical for neuronal integrity and functioning. However, the precise mechanisms by which microtubule dynamics are regulated during the development of the nervous system remain poorly understood. Here, we use global phosphoproteomic screening to identify cytoskeletal substrates of Ser-Arg Protein Kinase (SRPK), which is implicated in several neurodevelopmental and neurodegenerative conditions. We show that SRPK directly phosphorylates Microtubule Associated Protein (MAP)1S at multiple sites in a C-terminal region involved in proteolytic maturation and microtubule binding. SRPK-dependent MAP1S phosphorylation modulates the affinity of the MAP1S microtubule binding domain for microtubules and MAP1S proteolytic processing by the Calpain (CAPN)10 protease. Finally, we show that MAP1S proteolytic processing occurs progressively during neurodevelopment via a specific CAPN10 expression switch, corresponding with MAP1S acquisition of microtubule binding activity. Our results demonstrate a role for SRPK in coordinating processing and functionalisation of a key microtubule regulatory protein during neurodevelopment and provide insight into mechanisms by which the microtubule cytoskeleton may be dysregulated in neurological diseases.

## INTRODUCTION

Ser-Arg protein kinases (SRPKs) specifically phosphorylate Ser-Arg dipeptides ^1^ frequently found in Ser-Arg-rich splicing factors (SRSFs), which promote spliceosome complex assembly and pre-mRNA splicing ^2,3^. There are three SRPK family members: SRPK1 ^2^, SRPK2 ^4^ and SRPK3 ^5^, which have distinct, tissue-specific expression patterns, with SRPK1 mainly expressed in the testis, SRPK2 in the brain, and SRPK3 largely muscle-specific ^3,5,6^. In addition to splicing regulation, SRPKs perform other non-splicing functions, such as chromatin reorganization ^7,8^, zygotic genome reprogramming ^9^, transcriptional regulation ^10^, cell cycle progression, apoptosis induction, pluripotency, and differentiation ^10–12^.

SRPK plays an emerging role in neuronal development and functioning. SRPK mutants in *Drosophila* show impairment in life expectancy and locomotor behaviour ^13^, whilst SRPK79D, a neuron-specific SRPK homolog in *Drosophila*, phosphorylates active zone scaffold proteins at synapses ^13,14^. Mammalian SRPK1 and SRPK2 have also been shown to phosphorylate scaffold proteins, suggesting an evolutionarily conserved mechanism controlling axonal transport of active zone scaffold components^13^. SRPK2 is expressed in the synapses of hippocampal neurons, where it regulates self-assembly of the scaffold protein CAST1/ERC2, and maintains homeostatic plasticity, synaptic transmission release and presynaptic assembly by phosphorylating Rab3-interacting molecule 1 (RIM1) ^15,16^.

As a result of its roles in neurons, SRPK is implicated in neurodevelopmental disorders. Point mutations in SRPK3 cause X-linked intellectual disability, whilst deletions in SRPK2 are associated with intellectual disability ^17–21^. Knockout of the SRPK3 orthologue in zebrafish causes cerebellar atrophy and abnormal eye movement, phenotypes that resemble clinical features found in human intellectual disability ^17^, whilst SRPK phosphorylates the E3 ubiquitin ligase RNF12/RLIM, which is mutated in the X-linked intellectual disability disorder Tonne-Kalscheuer syndrome (TOKAS) ^7,22,23^. Finally, truncating or missense mutations of SRPKs have also been identified in other neurodevelopmental disorders, such as autism ^24^.

The microtubule cytoskeleton plays a significant role in maintaining cellular structure, morphogenesis, facilitating intracellular trafficking, and in dynamic cellular processes such as neural migration or neurite formation during neuronal development ^25–29^. Microtubule dynamics are regulated by various accessory proteins, particularly Microtubule Associated Proteins (MAPs), which promote microtubule polymerization and stabilization in neurons ^30^. Dysfunction of MAPs via mutation, abnormal phosphorylation, or improper proteolytic processing, can lead to abnormal microtubule dynamics, disrupting cellular integrity, intracellular transport, axonal growth, and synaptic function ^31–37^. For example, SRPK contributes to hyperphosphorylation of Tau in neurodegenerative disorders, which results in reduced microtubule dynamics, formation of paired helical filaments, impaired axonal transportation, neurite integrity, and neuronal death, ultimately contributing to disease pathogenesis ^38–45^. MAP dysfunction can also underpin pathogenesis of neurodevelopmental disorders ^46–51^. For example, MAP1B dysfunction leads to impaired synaptic plasticity and neuronal growth, affecting cognitive development and causing intellectual disability ^49^, whilst MAP1S dysfunction is associated with CDKL5 deficiency disorder (CDD), a neurodevelopmental disease caused by *CDKL5* mutations ^47,52^. However, how exactly phosphorylation regulates microtubule dynamics during neuronal development and degeneration of the nervous system remains poorly understood.

Here, we use global phosphoproteomic screening to uncover MAP1S as a major novel SRPK substrate. SRPK directly phosphorylates MAP1S at multiple sites in a C-terminal region involved in proteolytic maturation and microtubule binding. We show that SRPK-dependent MAP1S phosphorylation modulates both proteolytic processing of MAP1S into heavy and light chain fragments by the Calpain (CAPN)10 protease and the affinity of the MAP1S light chain microtubule binding domain for microtubules. Finally, we show that CAPN10 expression and MAP1S proteolytic processing are concomitantly and progressively induced during neurodevelopment, which corresponds with MAP1S acquisition of microtubule binding activity. Our results demonstrate a key role for SRPK in controlling the processing and functionalisation of a key microtubule regulatory protein during neurodevelopment and provide insight into mechanisms by which the microtubule cytoskeleton may be dysregulated in neurological diseases.

## EXPERIMENTAL PROCEDURES

### Plasmid constructs and site-directed mutagenesis

Site-directed mutagenesis and plasmid preparation were carried out by MRC-PPU Reagents & Services (https://mrcppureagents.dundee.ac.uk). All final plasmids were verified by Sanger sequencing performed by the MRC-PPU Reagents & Services Sequencing Laboratory (https://dnaseq.co.uk). Mutations were introduced into the corresponding wild-type recombinant DNA constructs using either KOD Hot Start DNA Polymerase (Merck, 71086) for PCR-based site-directed mutagenesis or the NEBuilder® HiFi DNA Assembly Kit (New England Biolabs, E5520) for fragment assembly, following the manufacturers’ protocols. Primers for mutagenesis were designed using SnapGene software. For the KOD Hot Start method, the parental methylated plasmid templates were digested with DpnI at 37 °C for 30 minutes after PCR amplification, followed by heat inactivation at 80 °C for 20 minutes before transformation. For the NEBuilder HiFi method, the PCR-amplified vector backbone and the mutation-containing DNA fragment were assembled in a reaction incubated at 50 °C for 60 minutes before transformation.

Transformed DH5α cells (MRC-PPU Reagents & Services) were prepared by incubating 2 µL of the chilled assembled product, following the standard transformation protocol. Colonies were inoculated overnight in LB medium with ampicillin and then purified using the QIAprep spin miniprep kit (Qiagen, 27104). All introduced mutations were verified by DNA sequencing by MRC-PPU Reagents & Services (https://dnaseq.co.uk).

### Cell culture and transfection

Male mouse embryonic stem cells (mESCs) were obtained from the laboratory of Dr. Janet Rossant (SickKids Research Institute, Toronto). Wild-type (*Map1s*^+/+^) and MAP1S knockout (*Map1s*^-/-^) mESCs were maintained on 0.1% (w/v) gelatin-coated tissue culture plate in DMEM with 10% FBS (v/v), 5% knockout serum replacement (KSR) (v/v), 2 mM glutamine, 0.1 mM non-essential amino acids (NEAA), 1 mM sodium pyruvate, 50 U penicillin, 50 μg/mL streptomycin, 0.1 mM β-mercaptoethanol (BME) and 20 ng/mL GST-tagged Leukaemia inhibitory factor (GST-LIF, MRC-PPU Reagents & Services, DU1715) media (ES-DMEM media), and incubated at 37°C and 5% CO_2_. cDNA plasmids were transfected into mESCs using Lipofectamine LTX or Lipofectamine 2000 or 3000 (Thermo Fisher Scientific, 11668019 or L300015) according to the manufacturer’s instructions. Cells were then cultured for another 24 hours before analysis. All cells were tested monthly for mycoplasma contamination.

### Preparation of cell extracts

Cells were washed with PBS twice and lysate made in lysis buffer (20 mM Tris–HCl pH 7.4, 150 mM NaCl, 1 mM EDTA, 1% NP-40 [v/v], 0.5% sodium deoxycholate [w/v], 10 mM β-glycerophosphate, 10 mM sodium pyrophosphate, 1 mM NaF, 2 mM Na_3_VO_4_, and 0.1 U/mL Complete Protease Inhibitor Cocktail Tablets (Roche, 11697498001), 80-200 μL to a well in a 6-well plate, 500 μL for a 10 cm cell culture dish and cells were scraped from the plate surface using a cell scraper to collected in a 1.5 mL Eppendorf tube. Lysates were incubated on ice for 15 minutes, then centrifuged at 4°C for 15 minutes at maximum speed (17,000 × g) to clear the lysate. The supernatant was collected and either used on the day or stored at-80°C. BCA colorimetric assay kit (Thermo Fisher Scientific, A55860) was used to measure the protein concentration of cell lysates according to the manufacturer’s instructions. A serial gradient concentration of bovine serum albumin (BSA) protein (0, 125, 250, 500, 1000, and 2000 μg/mL) was used to generate a standard concentration curve after quantifying the absorbance at 562 nm of each standard sample. 3 μL of lysate was mixed with 150 μL of the BCA mix in a 96-well plate and incubated at 37°C for 20-30 minutes. The concentration of each sample was then calculated according to the standard curve.

### Immunoblotting and Phos-tag immunoblotting

Endogenous or recombinant protein samples were denatured by incubating with 3x LDS and RA buffer at 95°C for 5 minutes and then loaded directly on commercial NuPAGE 4–12% Bis–Tris SDS–PAGE (Sodium Dodecyl Sulfate-Polyacrylamide Gel Electrophoresis) gels (Thermo Fisher Scientific). For Phos-tag™ SDS-PAGE, denatured protein samples were supplemented with 10 mM MnCl_2_ and loaded on 6% or 8% SDS-PAGE gels containing 0.375 M Tris-HCl pH 8.8, 0.001% SDS, 0.1 mM or 0.16 mM MnCl_2_, and 50 μM or 80 μM Phos-tag. After electrophoresis, gels were washed 3 times with transfer buffer containing 20 mM EDTA and once with 1x transfer buffer (48 mM Tris-HCl pH 8.3, 39 mM Glycine, 20% Methanol) without EDTA for 10 minutes at room temperature.

Proteins were then transferred to polyvinylidene fluoride (PVDF) (Merck Millipore) and incubated with primary antibodies diluted in TBS-T (20 mM Tris-HCl pH 7.5, 150 mM NaCl supplemented with 0.2% [v/v] Tween-20 [Sigma-Aldrich]) containing 5% non-fat milk buffer (w/v) at 4°C overnight. Following 3 times 10-minute washes with 1x TBS-T, secondary antibodies were incubated for 1 hour, followed by a further 3 times 10-minute washes with 1x TBS-T. HRP-linked secondary antibodies were visualized by Enhanced Chemiluminescence (Immobilon Western Chemiluminescent HRP substrate, Millipore) or SuperSignal West Pico PLUS Chemiluminescent Substrate (ThermoFisher Scientific) using a Chemidoc system (Bio-Rad). All protein signals were quantified using Image Lab software (Bio-Rad).

### Immunofluorescence

Cells were maintained in 12-well or 6-well plates with a gelatin-coated coverslip. 24 hours post-transfection, cells were fixed with 4% Paraformaldehyde (PFA) (w/v) diluted in PBS for 20 minutes in the dark. Fixed cells were then washed 3 times with PBS and permeabilized in 0.5% Triton X-100 (w/v) diluted in PBS for 5 minutes at room temperature. After the permeabilization, cells were blocked with 5% donkey serum for 1 hour at 37°C in a benchtop mini oven or overnight at 4°C. Coverslips were then washed 3 times with PBS and incubated with primary antibodies diluted in 1% donkey serum for 2 hours at 37°C. Coverslips were washed 3 times in PBS, incubated for 1 hour with fluorophore-tagged secondary antibodies diluted in 1% donkey serum at 37°C, incubated in 0.1 µg/ml Hoechst DNA stain for 5 minutes at room temperature in the dark. After washing in PBS, coverslips were mounted on microscope slides using microscopy-grade mounting media and dried at room temperature in the dark for at least 24 hours before imaging. Images were acquired using Zeiss 880 Airyscan microscope and DeltaVision Elite microscope (63x oil immersion objective, 20x immersion objective) and processed using Fiji software and Omero online website.

### mESC neural differentiation

mESCs were aggregated into embryoid bodies (EBs) using an EB medium (DMEM supplemented with 10% FBS (v/v), 2 mM glutamine, 0.1 mM non-essential amino acids (NEAA), 1 mM sodium pyruvate, 100 U/mL penicillin, 50 μg/mL streptomycin, 0.1 mM β-mercaptoethanol, and 2 μg/mL puromycin) containing 5 µM retinoic acid for 8 days. EBs were then dissociated into single cells using 0.05% Trypsin/EDTA (Thermo Fisher Scientific) and dissociated cells were seeded at a density of 100,000 cells/cm² of a 0.1% gelatin-coated 6-well plate, which suspended in N2B27 media (fresh 50% DMEM F12 (v/v), 0.5% N2 supplement (v/v), 50% Neurobasal media (v/v), 1% B27 supplement (v/v), 0.875 μM β-mercaptoethanol, 100 U Penicillin/Streptomycin, 2 mM L-glutamine) to promote further differentiation to neural cells for the rest 7 days. 20 ng/µL Fibroblast Growth Factor (FGF) and 20 ng/µL Epidermal Growth Factor (EGF) were added to N2B27 media in each well for the first 3 days to support initial differentiation and cell survival. The medium was replaced daily during this period to provide fresh nutrients and growth factors. After the initial 3 days, cells were maintained in only N2B27 medium without additional growth factors and maintained for 4 days. The media was refreshed every 2 days. At specified time points, cells were collected for further analysis to evaluate differentiation efficiency and to characterize the resulting cell populations.

### Human induced pluripotent stem cell (hiPSC) cortical neuron differentiation

hiPSCs were plated in TESR media containing 30 ng/mL bFGF and 10 ng/mL Noggin, at 60,000 cells/cm² in 10 cm dishes. The next day (Day 0 of differentiation), neuroepithelial induction was initiated by switching to Stage 1 media (DMEM F12, 0.2 μM LDN193189, 10 μM SB431542, 5 μM XAV939) and from Day 0 to Day 5. On Day 6, cells were detached using Accutase (Thermo Fisher Scientific) and cortical neuronal precursors were generated by introducing Stage 2 media (DMEM F12, 0.2 μM LDN193189, 10 μM SB431542, 5 μM XAV939, 0.1 μM Retinoic acid, 10 μM Y27632) at a density of 100,000 cells/cm². The cells were fed daily with Stage 2 media. On Day 12, the cells were changed into Stage 3 media (DMEM F12, 2.5 μM DAPT, 0.1 mM db-cAMP, 0.5 μM Retinoic acid, 200 ng/μL Ascorbic acid, 10 ng/mL BDNF, 10 ng/mL GDNF, 10 μM Y27632) for terminal cortical neuron maturation. The final plating density was 150,000 cells/cm², with 2 mL media for 6-well plates, 7.5 mL for 10 cm plates, and 100 µL for 96-well plates. Cells were maintained in stage 3 media with media changed every 2-3 days until Day 20. At specified time points, cells were collected for further analysis to evaluate differentiation efficiency and to characterize the resulting cell populations.

### Generation of *Map1s*^-/-^ mESCs

WT mESCs were transfected with CRISPR/Cas9 vectors, consisting of a pBabeD P U6 vector expressing the MAP1S (mouse) ex5 KO sense guide RNA and puromycin resistance cassette for selection, together with a pX335 vector expressing MAP1S (mouse) ex5 KO antisense guide RNA and Cas9 nickase (Cas9 D10A) enzyme. After 24 hours of incubation (37°C, 5% CO₂), transfected cells were selected with 3 µg/mL puromycin for 48 hours, and single cells were sorted using FACS. Sorted cells were maintained in a 96-well plate to grow colonies until confluency (7 to 14 days). After confluency, cells were expanded and screened by western blotting for MAP1S to identify putative *Map1s*^-/-^ mESC clones. Further confirmation was obtained by genomic DNA sequencing.

### Protein purification

GST-tagged proteins were expressed from pGEX6P-1 vectors in BL21 Codon Plus cells. A starter culture was used to inoculate 1 L LB media containing carbenicillin, grown at 37 °C with shaking at 200 rpm to an OD_600_ of 0.8. The temperature was then lowered to 15 °C and expression was induced with 0.01 mM Isopropyl β-D-1-thiogalactopyranoside (IPTG) for 18 hours at 200 rpm. Cells were harvested by centrifugation at 4000 × g for 30 minutes and resuspended in 20 mL ice-cold lysis buffer (50 mM Tris-HCl pH 7.5, 250 mM NaCl, 1% Triton X-100, 1 mM EDTA, 1 mM EGTA, 0.1% β-mercaptoethanol, 0.2 mM PMSF, 1 mM benzamidine). Cells were lysed by brief freeze–thawing at −80 °C followed by sonication (Branson Digital Sonifier) using 10x 15-second pulses at 45% amplitude. The lysate was clarified by centrifugation at 30,000 × g for 30 minutes at 4 °C.

The supernatant was incubated with 2 mL GSH–agarose beads (Abcam), pre-equilibrated in 50 mM Tris-HCl pH 7.5, for 1 hour at 4 °C with gentle rotation. Beads were washed in wash buffer (50 mM Tris-HCl pH 7.5, 250 mM NaCl, 0.1 mM EGTA, 0.1% β-mercaptoethanol) and bound proteins were eluted in the same buffer supplemented with 20 mM reduced glutathione (pH 7.5). Eluates were dialysed overnight at 4 °C into storage buffer (50 mM Tris-HCl pH 7.5, 0.1 mM EGTA, 150 mM NaCl, 0.5 mM TCEP, 270 mM sucrose) and stored at −70 °C.

### *In vitro* kinase assay

0.1 µg recombinant SRPK2 wild-type or kinase-inactive (SRPK2 D541A) were incubated with 0.5 µg GST-tagged human MAP1S fragments in kinase buffer (50 mM Tris-HCl pH 7.5, 0.1 mM EGTA, 10 mM MgCl_2_, 2 mM DTT) containing 2 mM ATP. Reactions were incubated at 30 °C for 30 minutes and stopped by adding 3x LDS/RA protein denaturation mix followed by boiling for 5 minutes.

For *in vitro* kinase assays with the SRPK inhibitor-resistant mutant (SRPK2 F164E), 500 ng C-terminal HA-tagged SRPK2 F164E and SRPK2 WT were transfected into WT mESCs. Cells were treated with or without SRPKIN-1 inhibitor (5 µM) before lysis. 500 µg of protein lysate from SRPK2 WT-and SRPK2 F164E-transfected cells were subjected to immunoprecipitation with anti-HA beads to purify SRPK2 WT and SRPK2 F164E, respectively. The beads were then used for the subsequent *in vitro* kinase assay by incubating with 0.5 µg recombinant mouse RNF12 substrate at 30 °C for 30 minutes. Reactions were stopped by adding 3x LDS/RA protein denaturation mix and boiling for 5 minutes.

For *in vitro* thiophosphorylation assay, an ATP analogue (ATPγS) was used instead of ATP, and the reaction was quenched using 20 mM EDTA. Thiophosphorylated proteins were then alkylated with 2.5 mM para-nitrobenzyl mesylate (PNBM), and the mixture was incubated at room temperature for 1 hour. Finally, the reaction was terminated by boiling for 5 minutes in a 3x LDA/RA buffer before proceeding with immunoblotting.

### Microtubule co-sedimentation assay

Taxol-stabilized microtubules were prepared by incubating 20 µM purified tubulin protein with 1 mM GTP and 1 mM DTT in BRB80 buffer (80 mM PIPES pH 6.8, 1 mM MgCl_2_, 1 mM EGTA) for 5 minutes at 4 °C. The mixture was then warmed to 37 °C, and Taxol was added stepwise in 1/100 volumes to a final concentration of 20 µM, 200 µM, and 2 mM, with each addition followed by a 5-minute incubation at 37 °C, except for the final step, which was incubated for 15 minutes. After stabilization, 10 µM microtubules were incubated with 3 µM MAP1S for 20 minutes at 25 °C. A 10 µL sample of the protein mixture was set aside as the input, while the remaining mixture was layered on top of a 40% glycerol cushion in an ultracentrifuge tube. The tubes were then ultracentrifuged for 30 minutes at 25 °C. Following centrifugation, 25 µL of the supernatant was collected, the pellet was washed 3 times with BRB80 buffer and resuspended in 25 µL of BRB80 supplemented with 20 µM Taxol before 25% of each fraction was analysed by immunoblotting.

To investigate the regulation of phosphorylation in microtubule binding, phosphorylation assays were conducted following the *in vitro* kinase assay method using 2 µg of wild-type and kinase-dead SRPK2 prior to the co-sedimentation assay. In the subsequent co-sedimentation assay, 11 µg of GST-MAP1S protein fragments were used as substrates at a concentration of 3 µM in the co-sedimentation assay.

### Phosphoproteomic profiling

Full details of this protocol have been published previously ^53^. In brief, mESCs were cultured in LIF/FBS as described and treated with 2 µM or 10 µM SRPKIN-1. Cells were then lysed, and proteins extracted and digested with trypsin. Phosphopeptides were enriched by using TiO_2_ microspheres (GL Sciences), followed by TMT labelling (Thermo Fisher Scientific). The samples were then fractionated through basic C18 reverse-phase chromatography (bRP) and analysed by LC-MS/MS. The LC-MS/MS data were processed using Proteome Discoverer software v2.2 (Thermo Fisher Scientific). To identify peptides that were significantly modified after 5 and 20 minutes of stimulation, the limma package (Bioconductor) was utilized.

### Targeted phosphosite mapping

For phospho-site identification, protein samples were analysed by SDS-PAGE and stained with InstantBlue® Coomassie Protein Stain (ThermoFisher Scientific). The target bands were excised and dissected into 1 mm^2^ cubes and then washed with water, 50% acetonitrile (ACN)/water, 0.1 M NH_4_HCO_3_, and 50% ACN/50 mM NH_4_HCO_3_ (All from Sigma-Aldrich). Every wash was 0.5 mL for each band and lasted for 10 minutes. After washing, protein cubes were incubated in 10 mM DTT/0.1 M NH_4_HCO_3_ at 37 °C for 20 minutes and alkylated in 50 mM iodoacetamide/0.1 M NH_4_HCO_3_ in the dark for 20 minutes. Gel cubes were destained in 50 mM NH_4_HCO_3_ and 50% ACN/50 mM NH_4_HCO_3_, then shrunk with 0.3 mL acetonitrile for 15 minutes. Gel cubes were dried in a Speed-Vac and swelled in 25 mM Triethylammonium bicarbonate containing 5 µg/mL of Trypsin and shaken at 30 °C overnight for proteolytic digestion. 0.3 mL acetonitrile was added for washing for 15 minutes, then the supernatant was transferred to a clean tube and frozen at-80 °C before being dried using Speed-Vac. Then samples were sent for mass spectrometry analysis.

For liquid chromatography followed by mass spectrometry (LC-MS), an Orbitrap Exploris 240 coupled to an Evosep One nano-LC system was used to acquire mass spectrometry data in data-dependent mode using the following parameters: MS1 spectra were acquired at a resolution of 60,000 (at 400 m/z) for a mass range of 375-1500 m/z with an FTMS full AGC target of 300%. The most intense ions (with a minimal signal threshold of 5000) were selected for MS2 analysis at a resolution of 15000 with a standard AGC target and were fragmented using a Higher Collision Dissociation (HCD) collision energy of 30% and then dynamically excluded for 30 seconds.

Data was analysed using Proteome Discoverer v.2.4 and Mascot using MRC_Database_1 (2,110 sequences; 1,374,911 residues). Parameters used were the following: Variable modifications: Oxidation (M), Dioxidation (M), Phospho (STY); Fixed modifications: Carbamidomethyl (C), Enzyme: Trypsin/P, Maximum missed cleavages: 2, Precursor tolerance: 10ppm, MS2 tolerance: 0.06Da, Minimum score pep-tides: 17. Phospho-site assignment probability was estimated via Mascot and IMP-ptmRS (Proteome Discoverer v.2.4).

### Antibodies

In-house antibodies were generated by MRC-PPU Reagents & Services (https://mrcppureagents.dundee.ac.uk). For immunoblotting, antibody solutions were diluted in 5% (w/v) skimmed milk in TBS-T or 5% (w/v) bovine serum albumin (BSA) in TBS-T.

Antibodies used were as follows: anti-mouse IgG, HRP (Cell Signaling Technology, 7076S), anti-rabbit IgG, HRP (Cell Signaling Technology, 7074S), and anti-sheep IgG, HRP (MRC PPU Reagents and Services, DU-SPI116). MAP1S antibodies were from Thermo Fisher Scientific (PA5-100607), Sigma-Aldrich (HPA050934), and Proteintech (15695-1-AP). Phospho-specific MAP1S antibodies were from MRC-PPU Reagents & Services (DU-SA385 for S812, DU-SA339 for S900 ^52^, and DU-DA213 for S786). Tag antibodies were anti-HA, HRP (Roche, 12013819001), anti-HA (Abcam, ab9110), anti-Flag (Sigma-Aldrich, F1804-50), anti-Flag, HRP (Sigma-Aldrich, A8592-2), and anti-GST (MRC PPU Reagents and Services, DU-S902A). Additional antibodies were anti-thiophosphate ester (Abcam, ab92570), anti-ERK1/2 (MRC-PPU Reagents & Services, DU-S221B), anti-β-actin (Cell Signaling Technology, 4967), anti-α-tubulin (Proteintech, 66031-1-Ig), and anti-MAP2 (Sigma-Aldrich, M1406-100UL). Stem cell and neuronal markers included anti-OCT4 (Abcam, ab19857), anti-NANOG (Cell Signaling Technology, 3580S), anti-βIII-tubulin (Novus Biologicals, NB100-1612), anti-PAX6 (BioLegend, 901302), anti-SOX2 (Cell Signaling Technology, 4900S), anti-FOXG1 (Abcam, ab196868), and anti-TBR1 (Abcam, ab31940). For immunofluorescence, Alexa Fluor–conjugated secondary antibodies were from Thermo Fisher Scientific, including rabbit IgG Alexa Fluor® 488 (A21206), chicken IgG Alexa Fluor® 488 (A11039), mouse IgG Alexa Fluor® 647 (A31571), and rabbit IgG Alexa Fluor® 594 (A21207).

### Data analysis

Each experiment was performed at least 3 times. For statistical analysis, at least 3 biological replicates were used, with individual points representing a single biological replicate. XIC (extracted ion chromatogram) mass spectrometry data were obtained using Freestyle software (Thermo Fisher Scientific). Paired and unpaired Student’s t-tests were performed for XIC mass spectrometry data analysis using GraphPad Prism V10.00 software. Statistical significance was indicated as follows: not significant (p > 0.05), significant (*, p < 0.05). Basic data analysis for mean differences was performed using Excel software. Immunoblots were quantified by densitometric analysis in Image Lab software (BioRad), with data presented as the mean ± SEM of at least 3 biological replicates.

## RESULTS

### Global phosphoproteomics identifies putative Ser-Arg Protein Kinase (SRPK) substrates

In order to identify novel SRPK substrates, we performed a global phosphoproteomic screen in mouse embryonic stem cells (mESCs) following treatment with 2 µM or 10 µM of the potent and selective SRPK inhibitor SRPKIN-1 ^54^ (Fig. 1A). As SRPKIN-1 is a covalent inhibitor, we used a ‘wash-out’ strategy in which mESCs are incubated with SRPKIN-1 for 3 hours followed by media change and further 2 hours incubation to increase selectivity ^54^. This analysis identified a total of 29,324 phosphopeptides from 3 biological replicates, including cohorts of phosphopeptides that are significantly reduced in abundance upon SRPKIN-1 treatment (Fig. 1B). Most reduced abundance phosphopeptides identified upon 2 µM SRPKIN-1 treatment are also identified at 10 µM (Fig. 1C). See (https://shiny.compbio.dundee.ac.uk/mgierlinski/private/phospho_srpk/pep/) for an interactive data explorer and (https://www.compbio.dundee.ac.uk/user/mgierlinski/temp/phospho_srpk/report.html) for a full summary of the data and analysis.

**Figure 1:**
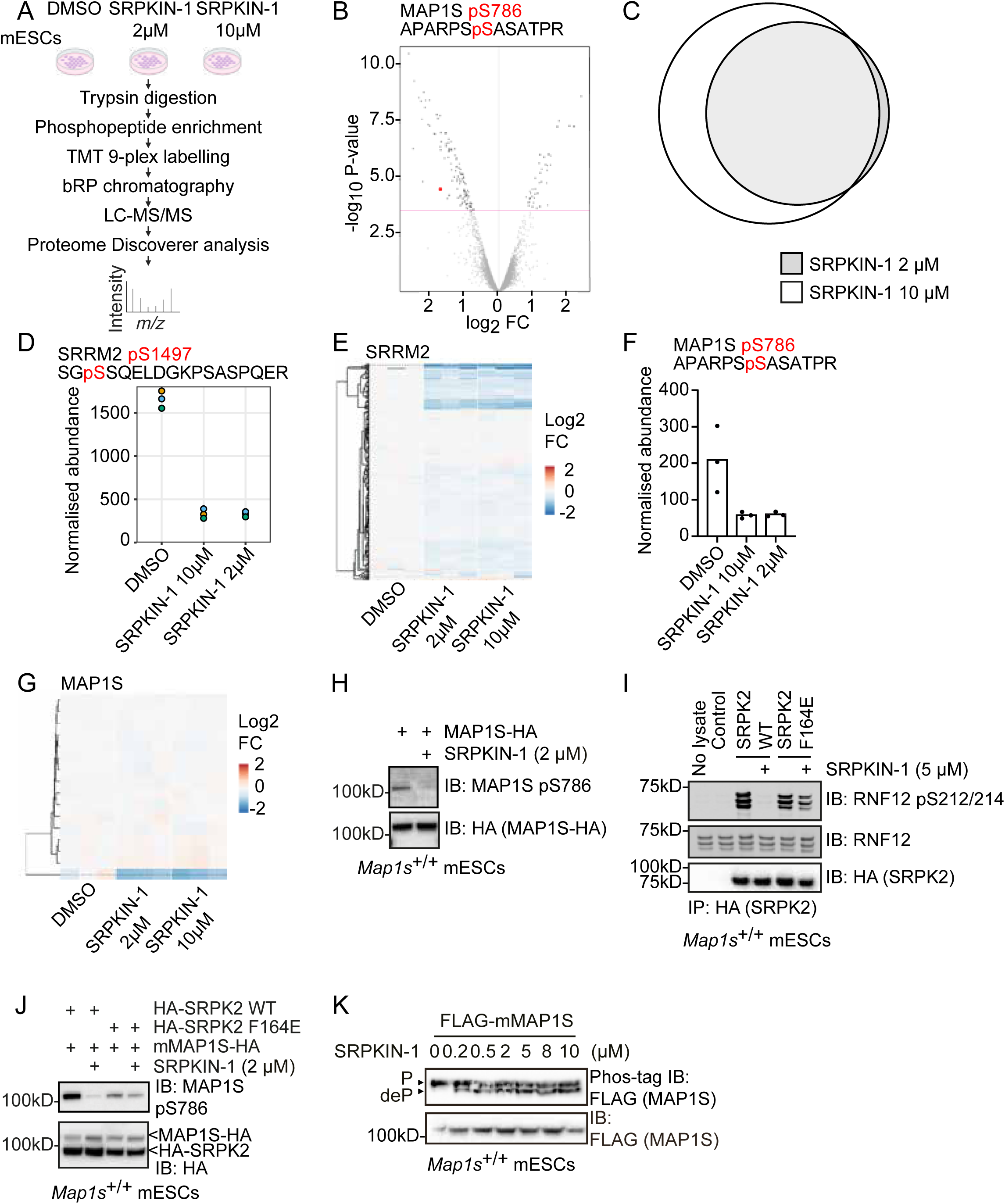
Global phosphoproteomics identifies MAP1S as a potential SRPK substrate (A) Phosphoproteomic workflow; mouse embryonic stem cells (mESCs) were treated with DMSO, or 2 µM or 10 µM SRPKIN-1 for 3 hours, followed by 2 hours washout. Proteins were extracted, trypsin digested, and phosphopeptides enriched and labelled with Tandem Mass Tags (TMT) for multiplexed quantification. The labelled peptides underwent reverse-phase (bRP) chromatography before Liquid Chromatography-Tandem Mass Spectrometry (LC-MS/MS) analysis. The resulting data were processed and analysed using Proteome Discoverer software (n=3). The figure was generated using BioRender. (B) Volcano plot of phosphopeptide abundance in DMSO control compared to 2 µM SRPKIN-1. The Y-axis value above the red line corresponds to False Discovery Rate (FDR) <0.05; FC represents the fold change in phosphopeptide abundance. Negative log_2_ FC values indicate a decrease in phosphorylation abundance upon SRPKIN-1 treatment (n=3) (C) Venn diagram showing overlap of phosphopeptides showing significant change in abundance following 2 µM and 10 µM SRPKIN-1 treatment. (D) Phosphopeptide abundance of SRRM2 pS1497 phosphopeptide (SGpSSQELDGKPSASPQER) upon DMSO, 2 µM, and 10 µM SRPKIN-1 treatment (n=3). (E) Hierarchical cluster showing change in abundance of all SRRM2 phosphopeptides identified by phosphoproteomics. (F) Phosphopeptide abundance of MAP1S pS786 phosphopeptide (APARPpSSASATPR) upon DMSO, 2 µM, and 10 µM SRPKIN-1 treatment (n=3). (G) Hierarchical cluster showing change in abundance of all MAP1S phosphopeptides identified by phosphoproteomics (n=3). (H) WT (*Map1s*^+/+^) mESCs were transfected with mMAP1S-HA and treated with 2 µM SRPKIN-1 for 6 hours, followed by immunoblotting (IB) with anti-MAP1S phospho-S786 and anti-HA antibodies (n=3). (I) WT (*Map1s*^+/+^) mESCs were transfected with HA-SRPK2 WT, HA-SRPK2 F164E, or EV control. HA-SRPK2 was immunoprecipitated and incubated with recombinant RNF12 in a kinase assay, followed by immunoblotting (IB) with anti-pS212/214 RNF12, anti-RNF12, and anti-HA antibodies (n=3). (J) mouse MAP1S-HA were either co-transfected with HA-SRPK2 WT or HA-SRPK2 F164E or EV control into WT (*Map1s*^+/+^) mESCs and treated with either DMSO or 2µM SRPKIN-1 for 6 hours. Results were analysed by immunoblotting (IB) with anti-MAP1S phospho-S786, anti-MAP1S, and anti-HA antibodies (n=3). (K) WT (*Map1s*^+/+^) mESCs were transfected with FLAG-mMAP1S, treated with increasing concentrations of SRPKIN-1, and analysed by either phos-tag (top) or regular (bottom panel) immunoblotting (IB) for FLAG. P = phosphorylated MAP1S; deP = dephosphorylated MAP1S (n≥3).

To provide proof of concept support for this analysis, we first hypothesised that the cohort of phosphopeptides downregulated upon SRPK inhibition is likely to include known SRPK substrates. Indeed, phosphopeptides from expected SRPK substrates such as the splicing factor Ser/Arg repetitive matrix protein (SRRM)2 are identified using this workflow (Fig. 1D, E). Many other potential SRPK substrates are also identified, including cytoskeleton regulatory components. Of particular interest, microtubule-associated protein (MAP)1S, a member of a family of microtubule regulatory proteins implicated in neurological disorders including neurodegeneration ^55^ and intellectual disability ^47^, is one of the major hits in this screen. A MAP1S S786 phosphopeptide is reduced in abundance upon SRPK inhibition with SRPKIN-1 (Fig. 1B). Individual data points for MAP1S S786 phosphopeptide upon SRPK inhibition at both 2 µM and 10 µM SRPKIN-1 are provided (Fig. 1F). Interestingly, 17 further MAP1S phosphopeptides were identified in this analysis, none of which are significantly impacted by SRPK inhibition (Fig. 1G).

### SRPK activity is required for Microtubule Associated Protein (MAP)1S phosphorylation at S786

MAP1S is a microtubule-binding protein comprising microtubule and actin-binding regions ^56,57^, which is cleaved by the Calpain (CAPN)10 protease between C841 and M842 in human MAP1S (C754 and M755 in mouse MAP1S) into heavy chain (HC) and light chain (LC) fragments ^58,59^. Our phosphoproteomics screen indicates that SRPK activity is required for mouse MAP1S phosphorylation at S786, which is within the microtubule-binding region of the LC. We therefore sought to validate the role of SRPK in regulating MAP1S phosphorylation by raising a phospho-specific antibody to MAP1S phospho-S786 to easily and robustly monitor MAP1S S786 phosphorylation. First, we expressed C-terminally HA-tagged mouse MAP1S (MAP1S-HA) constructs in mESCs and investigated MAP1S S786 phosphorylation. A strong signal is detected using the anti-MAP1S pS786 antibody with the expression of MAP1S-HA (Fig. 1H). However, this signal is abolished by treatment with the SRPK inhibitor SRPKIN-1 (Fig. 1H), providing orthogonal validation of our phosphoproteomic results, which indicate that MAP1S S786 is phosphorylated in an SRPK-activity-dependent manner (Fig. 1B).

### An inhibitor-resistant SRPK rescues MAP1S S786 phosphorylation upon SRPK inhibition

To further confirm the role of SRPK activity in modulating MAP1S S786 phosphorylation, and to rule out potential off-target kinases inhibited by SRPKIN-1, we sought to develop an inhibitor-resistant allele of SRPK. Using a series of mutations identified as driving kinase inhibitor resistance in cancer ^60^, we tested whether orthologous mutations render SRPK insensitive to SRPKIN-1. Using *in vitro* SRPK phosphorylation of the E3 ubiquitin ligase RNF12/RLIM ^21^ as a readout, we show that a F164E substitution in the gatekeeper pocket renders SRPK2 partially resistant to SRPKIN-1 inhibition, although SRPK2 F164E has slightly reduced kinase activity compared to SRPK2 WT (Fig. 1I).

We hypothesised that if SRPK is the MAP1S S786 kinase, an inhibitor-resistant SRPK will rescue MAP1S S786 phosphorylation even upon SRPK inhibition with SRPKIN-1. Therefore, we used SRPK2 F164E as an inhibitor-resistant mutant to determine whether MAP1S S786 phosphorylation is maintained in the face of SRPKIN-1 treatment. In the presence of SRPK2 WT, MAP1S S786 phosphorylation is abolished upon SRPKIN-1 treatment (Fig. 1J). In contrast, in the presence of SRPK2 F164E, MAP1S S786 phosphorylation is maintained at levels similar to those observed in the absence of SRPKIN-1 treatment (Fig. 1J). These data therefore suggest that inhibition of MAP1S S786 phosphorylation by SRPKIN-1 is via on-target inhibition of SRPK, and not via off-target inhibition of other kinases by SRPKIN-1.

### SRPK activity is required for Microtubule Associated Protein (MAP)1S phosphorylation at multiple motifs

In our phosphoproteomics analysis, a single MAP1S phosphorylation site was identified as significantly reduced upon SRPKIN-1 treatment, although other MAP1S phosphorylation sites were detected (Fig. 1; Table 1). Therefore, we sought to investigate the extent to which SRPK activity contributes to MAP1S phosphorylation. To this end, we employed Phos-tag SDS-PAGE analysis to visualise distinct MAP1S phosphorylated species. MAP1S resolves as a single phosphorylated species on Phos-tag (Fig. 1K). SRPK inhibition leads to the appearance of a less phosphorylated MAP1S species (Fig. 1K), even at concentrations of SRPKIN-1 down to 0.2 µM. Taken together, these results confirm that SRPK activity modulates MAP1S phosphorylation. Our Phos-tag SDS-PAGE results suggest that MAP1S sites in addition to S786, may be phosphorylated in an SRPK-dependent manner. To fully address this possibility, we first performed targeted phosphoproteomics to directly map all MAP1S phosphorylation sites in mESCs. Following expression of N-terminally FLAG-tagged mouse MAP1S (Flag-MAP1S) and SRPKIN-1 treatment, MAP1S was immunoprecipitated, analysed by SDS-PAGE and Coomassie staining, and MAP1S excised and analysed by LC-MS-MS (Fig. 2A). From 3 biological replicates, 16 MAP1S phosphopeptides were identified, comprising a total of 15 high-confidence phosphorylation sites. A summary of all MAP1S phosphopeptides identified is provided (Table 2).

**Figure 2:**
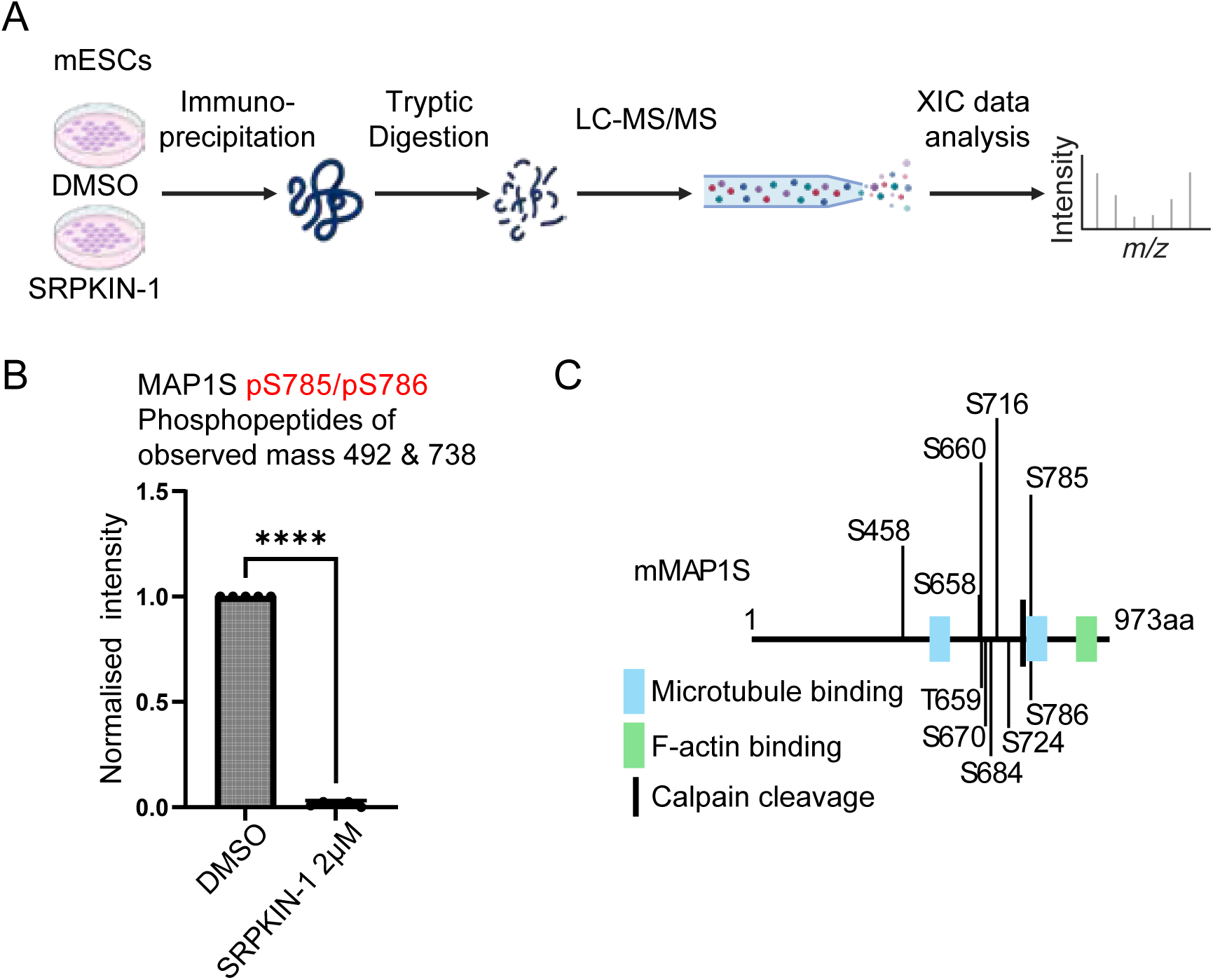
Mass Spectrometry identifies multiple SRPK-dependent phosphorylation sites on MAP1S C-terminus in mESCs (A) Schematic of targeted phosphopeptide mapping of SRPK-dependent MAP1S phosphorylation sites in mESCs. *Map1s^+/+^* mESCs were transfected with FLAG-mMAP1S and treated with DMSO or 10 µM SRPKIN-1 for 4 hours, followed by anti-FLAG immunoprecipitation of FLAG-MAP1S and analysis by LC-MS/MS (n=3). The figure was generated using BioRender. (B) Statistical analysis of Extracted Ion Chromatogram (XIC) values for abundance of MAP1S S785/785 phosphopeptides following DMSO and SRPKIN-1 treatment. Data are presented as mean ± standard deviation (SD) for identified peptides of the indicated observed mass that contain the indicated phosphorylation sites. Statistical significance was determined by the Shapiro-Wilk test for normal distribution, and a paired T-test was used for the group comparison on GraphPad Prism 9.4.0. **** = p<0.0001. (C) Diagram of mouse MAP1S (mMAP1S) showing the positions of MAP1S phosphorylation sites that are significantly decreased following SRPK inhibition. The Calpain (CAPN)10 cleavage site is located between C754 and M755. Blue = microtubule-binding region (aa755-875 in the light chain ^56^, specific location in the heavy chain not defined ^57^, green = actin-binding domain (aa852-973) ^56^.

**Table 1:** All MAP1S phosphopetides identified by global phosphoproteomics. The table shows all detected MAP1S phosphopeptides by global phosphoproteomics. High confidence assignment of phosphorylation sites is marked in red, lower confidence sites are marked in green.indicates ambiguity about which residue is phosphorylated. FC = fold change in phosphopeptide abundance between control (DMSO) and SRPK inhibitor (SRPKIN-1) treated groups. Negative log_2_FC indicates decreased abundance, and positive log_2_FC indicates increased abundance upon SRPK inhibition. False discovery rate (FDR) <0.05 is considered statistically significant.

| SEQUENCE | RESIDUES | Log<sub>2</sub>FC | FC | FDR |
| --- | --- | --- | --- | --- |
| APARPSASATPR | S786 | -1.7 | 0.308 | 0.00859 |
| STSPHDVDLCLVSPCEFSHR | S670, T659/S? | -0.156 | 0.898 | 1 |
| STSPHDVDLCLVSPCEFSHR | T659/S? | -0.127 | 0.916 | 1 |
| SQERPPETPPTSVSESLPTLSDSDPVP<br>VADSDDDAGSESAAR | S709, S724 | -0.193 | 0.875 | 1 |
| STSPHDVDLCLVSPCEFSHR | S670, S/T659? | -0.057 | 0.961 | 1 |
| RSTSPHDVDLCLVSPCEFSHR | T659, S660,<br>S670 | -0.12 | 0.920 | 1 |
| RSTSPHDVDLCLVSPCEFSHR | S658, T659/S? | -0.0208 | 0.986 | 1 |
| KPPPPASPGSSDSSAR | S684, S? | 0.191 | 1.142 | 1 |
| SGPGVVNTK | S520 | 0.0828 | 1.059 | 1 |
| ANSQDSLASR | S462 | -0.0281 | 0.981 | 1 |
| SQERPPETPPTSVSESLPTLSDSDPVP<br>VADSDDDAGSESAAR | S724 | 0.0385 | 1.027 | 1 |
| RPVVTTHDLEAPSRANSQDSLASR | S462 | 0.056 | 1.040 | 1 |
| RSTSPHDVDLCLVSPCEFSHR | T659/S? | -0.076 | 0.949 | 1 |
| SGPGVVNTKPR | S520 | 0.163 | 1.120 | 1 |
| KPPPPASPGSSDSSAR | S684 | 0.1 | 1.072 | 1 |
| ANSQDSLASRDSARK | S471 | 0.281 | 1.215 | 1 |

**Table 2:**
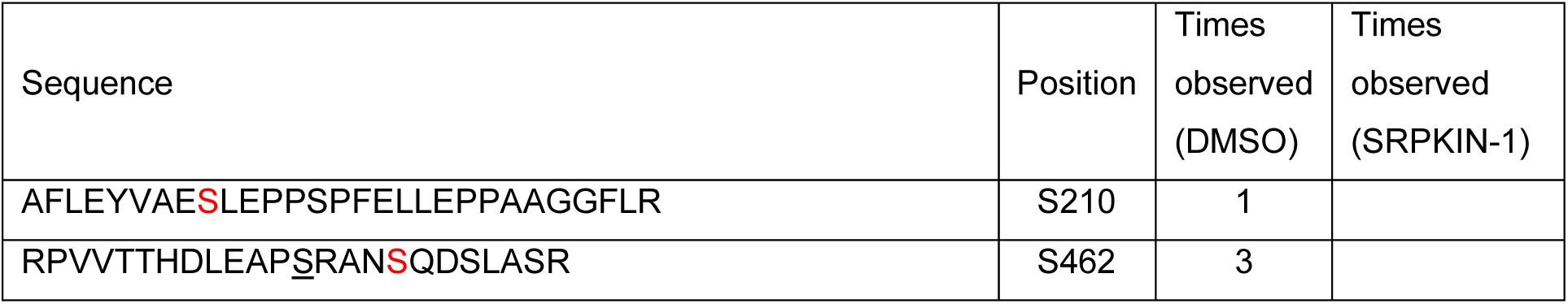
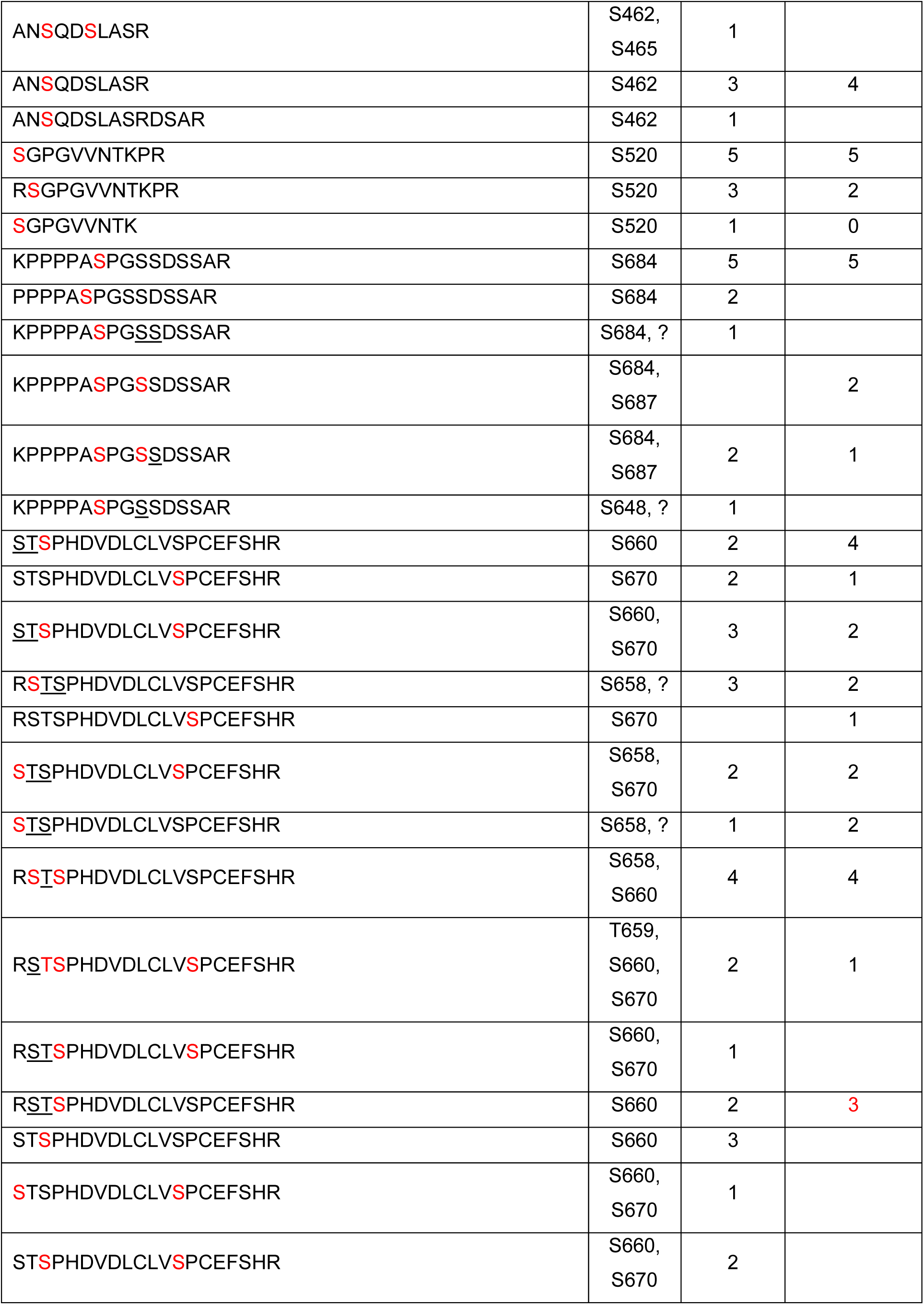
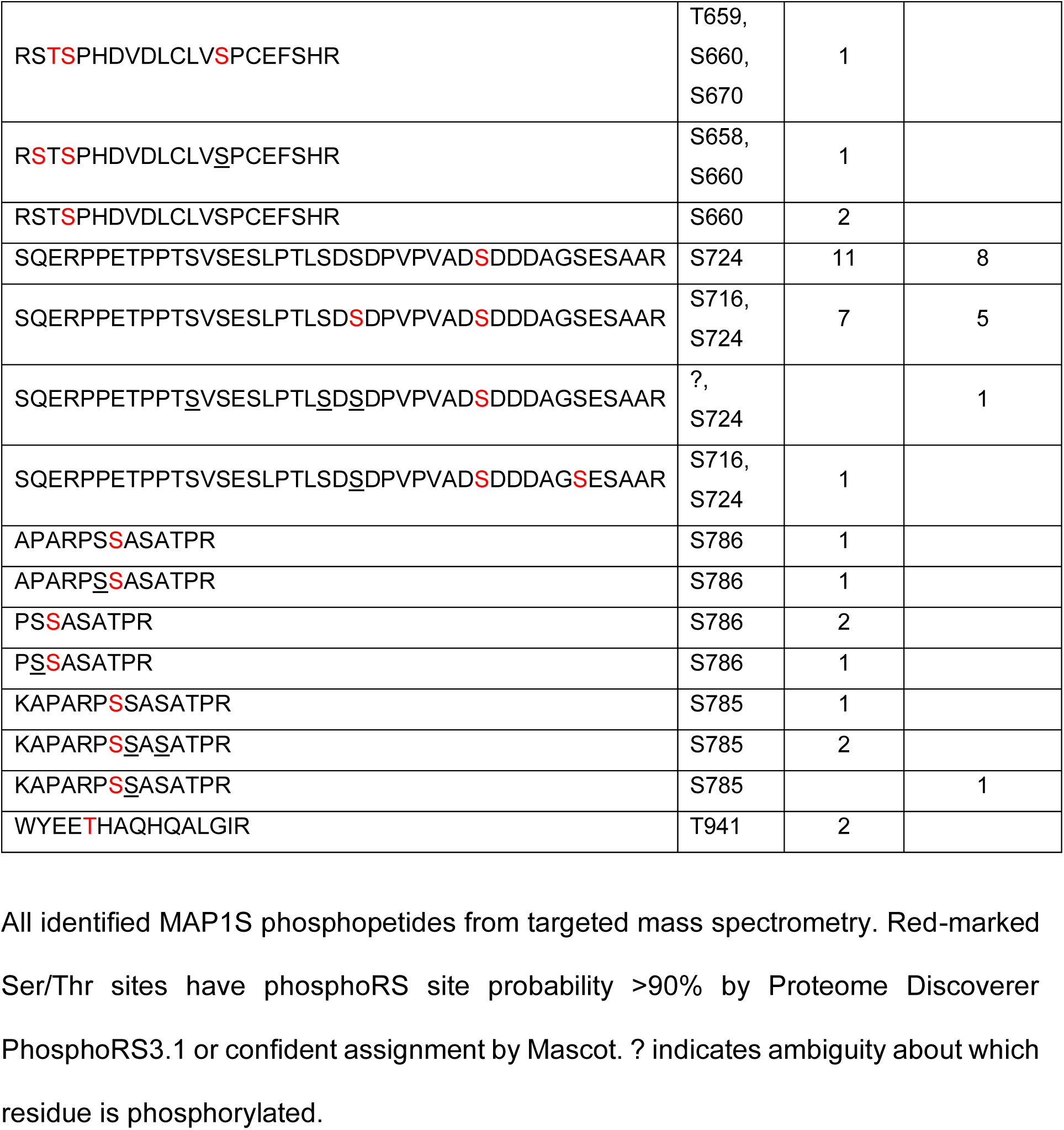
MAP1S phosphopeptides identified from mESCs with phosphoRS site probability >90%.

We next sought to quantify MAP1S phosphopeptide abundance to identify phosphorylation sites that are impacted by SRPK inhibition. As proof of principle, we show that a MAP1S phosphopeptide phosphorylated at S785 and S786 (S785/S786), of which S786 is identified in our global phosphoproteomic screen (Fig. 1), is significantly reduced in abundance following SRPKIN-1 treatment (Fig. 2B). In addition to MAP1S S785/S786, several other MAP1S phosphopeptides are significantly reduced in abundance following SRPKIN-1 treatment, including S658/T659/S660/S670, S716/S724, and S684 (Fig. S1). In contrast, a MAP1S S520 phosphopeptide is not significantly reduced in abundance upon SRPKIN-1 treatment (Fig. S1). A summary of high-confidence SRPK-dependent MAP1S motifs is presented (Fig. 2C). Interestingly, these phosphorylation motifs mostly lie in the MAP1S C-terminal region, which includes a site for MAP1S proteolytic cleavage into heavy chain (HC) and light chain (LC) polypeptides by the Calpain family protease CAPN10 ^58,59^, and motifs implicated in mediating MAP1 family binding to microtubules and actin^56,57,61–65^.

### SRPK directly phosphorylates the MAP1S C-terminal region

We next tested whether MAP1S is a direct substrate of SRPK. To this end, we purified from *E. coli* 4 recombinant GST-tagged MAP1S fragments covering the entire length of the human MAP1S protein (Fig. 3A, B). We then employed a thio-phosphorylation assay, which, instead of regular ATP, uses ATPγS followed by click-chemistry mediated modification to enable visualisation of thio-phosphorylation by immunoblotting ^66^. Consistent with our targeted phosphoproteomic identification of SRPKIN-1 sensitive phosphorylation sites in mESCs, SRPK2 specifically phosphorylates human MAP1S in the C-terminal region, specifically on fragments aa501-750 and aa751-1059 (Fig. 3C). We used Phos-tag immunoblotting to investigate the stoichiometry of GST-MAP1S aa501-750 and aa751-1059 phosphorylation following incubation with either wildtype kinase active SRPK2 (SRPK2 KA) or a kinase inactive mutant (SRPK2 KI). These data show that both SRPK2 phosphorylated MAP1S fragments are phosphorylated on a Phos-tag gel, with MAP1S fragment aa501-750 phosphorylated to high stoichiometry (Fig. 3D). These data confirm that the MAP1S C-terminus is an SRPK substrate *in vitro*, suggesting that the SRPKIN-1 sensitive MAP1S phosphorylation sites identified in mESCs are directly phosphorylated by SRPK.

**Figure 3:**
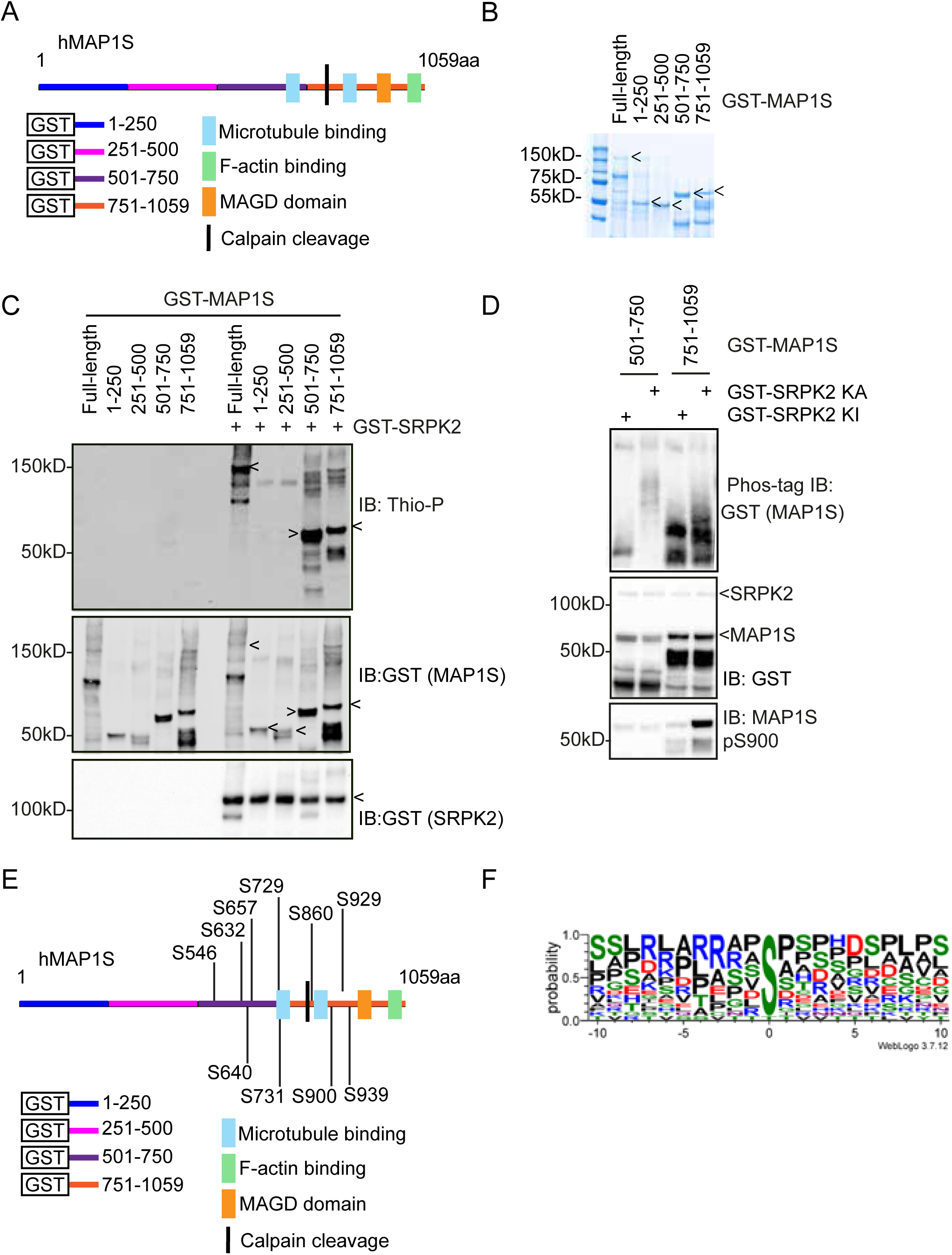
The MAP1S C-terminus is directly phosphorylated by SRPK in vitro (A) Diagram of human MAP1S (hMAP1S) protein structure with GST-MAP1S fragments aa1-250, aa251-500, aa501-750, and aa751-1059 highlighted. The Calpain (CAPN)10 cleavage site is located between C841 and M842. Blue = microtubule-binding region (aa867-945 in the light chain, location in the heavy chain not defined), yellow = mitochondrial accumulation and genome destruction (MAGD) domain (aa967-991), green = actin-binding domain (aa960-1059) ^76^. (B) Coomassie blue staining of purified hMAP1S full-length and fragments. (C) Active human GST-SRPK2 and GST-tagged hMAP1S fragments were incubated in an *in vitro* thiophosphate kinase assay and analysed by immunoblotting (IB) for anti-Thiophosphate (Thio-P) and anti-GST (n≥3). (D) Active human WT SRPK (GST-SRPK KA) or kinase inactive mutant SRPK (GST-SRPK KI) and indicated GST-tagged hMAP1S fragments were incubated in an *in vitro* thiophosphate kinase assay and analysed by regular immunoblotting (IB) with anti-GST (top) and anti-MAP1S pS900 antibodies (middle), and by Phos-tag immunoblotting (IB) with anti-GST antibodies (n=3). (E) Positions of SRPK phosphorylation sites identified on hMAP1S *in vitro*. (F) Sequence alignment of all identified SRPK-dependent phosphorylation sites on MAP1S. The X-axis is the amino acid position relative to the identified phosphorylation site (0 position). The height of symbols within the stack indicates the relative frequency of each amino acid. Figure generated using WebLogo (https://weblogo.threeplusone.com/).

Having shown that SRPK2 can directly phosphorylate recombinant human MAP1S C-terminal fragments *in vitro*, we next sought to identify the specific phosphorylation sites. We again performed targeted phosphoproteomics, this time on GST-MAP1S fragments aa501-750 and aa751-1059. Following phosphorylation by SRPK *in vitro*, GST-MAP1S aa501-750 and aa751-1059 were analysed by SDS-PAGE and Coomassie staining, and excised and analysed by LC/MS/MS. From 2 biological replicates, 12 phosphopeptides comprising a total of 12 phosphorylation sites were identified from GST-MAP1S aa501-750 and aa751-1059. A summary of all phosphopeptides identified is provided, along with total MAP1S coverage achieved (Fig. 3E & Table 3). Importantly, many of the direct SRPK phosphorylation sites identified on recombinant human MAP1S fragments correspond to phosphorylation motifs identified as sensitive to SRPK inhibition in mouse MAP1S in mESCs.

**Table 3:**
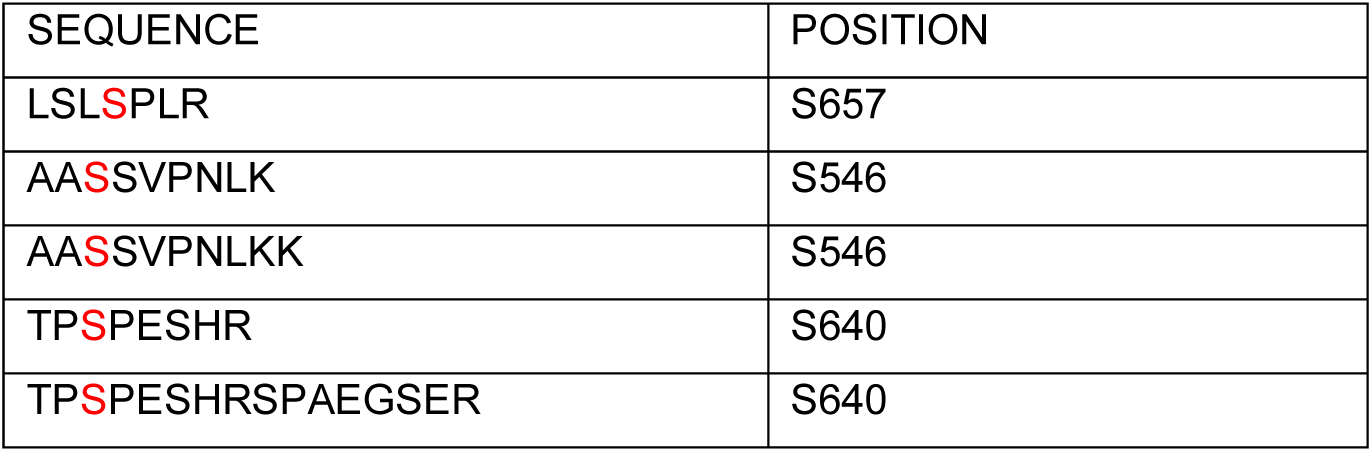
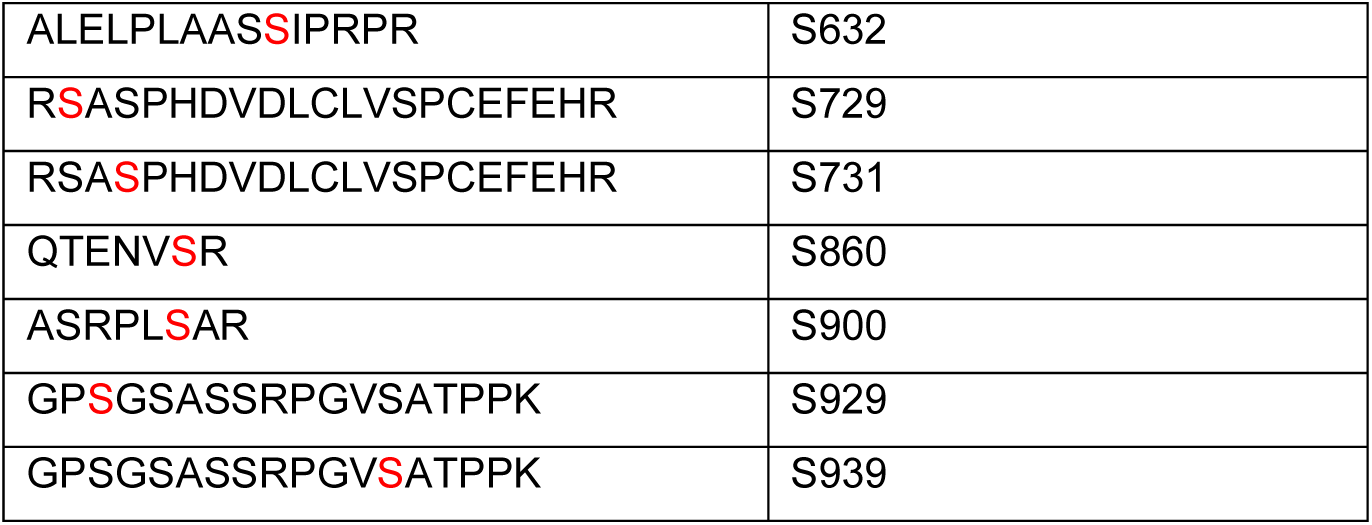
MAP1S phosphopeptides identified by mass spectrometry following *in vitro* phosphorylation by SRPK2. All identified phosphopetides from targeted mass spectrometry. Red-marked Ser/Thr sites have phosphoRS site probability >90% by Proteome Discoverer PhosphoRS3.1 or confident manual assignment from Mascot and MS2 data.

We further analysed the SRPK phosphorylation motifs identified on human and mouse MAP1S using WebLogo (https://weblogo.threeplusone.com/) to visualise motif enrichment around the identified phosphorylation sites. This analysis shows a preference for Arginine (R) at - 3 position and Proline (P) at + 1 position (Fig. 3F), consistent with the reported SRPK motif preference ^3,44,67,68^, suggesting that MAP1S phosphorylation by SRPK is conserved between mouse and human, despite the low overall sequence conservation of human and mouse MAP1S in the C-terminal region.

### SRPK phosphorylation inhibits MAP1S microtubule binding *in vitro*

We next sought to explore the function of MAP1S phosphorylation by SRPK. Previous studies have indicated that MAP1S phosphorylation in the C-terminal region proximal to the LC microtubule-binding motif can modulate interaction with microtubules ^47^. Therefore, we addressed the impact of MAP1S phosphorylation by SRPK on interaction with microtubules. We first tested whether recombinant GST-MAP1S fragments interact with purified polymerised microtubules using an *in vitro* co-sedimentation assay. As expected, microtubules sediment upon ultracentrifugation (Fig. 4A), whilst GST-MAP1S aa1-250 and aa251-500 remain mostly in the supernatant (Fig. 4A). However, GST-MAP1S aa751-1059, and to a lesser extent, GST-MAP1S aa501-750, co-sediment with microtubules (Fig. 4A). Importantly, none of the GST MAP1S fragments sediment upon ultracentrifugation in the absence of microtubules (Fig. 4B). These data confirm previous literature that MAP1S aa751-1059 encodes a major microtubule binding region ^56,57^.

**Figure 4:**
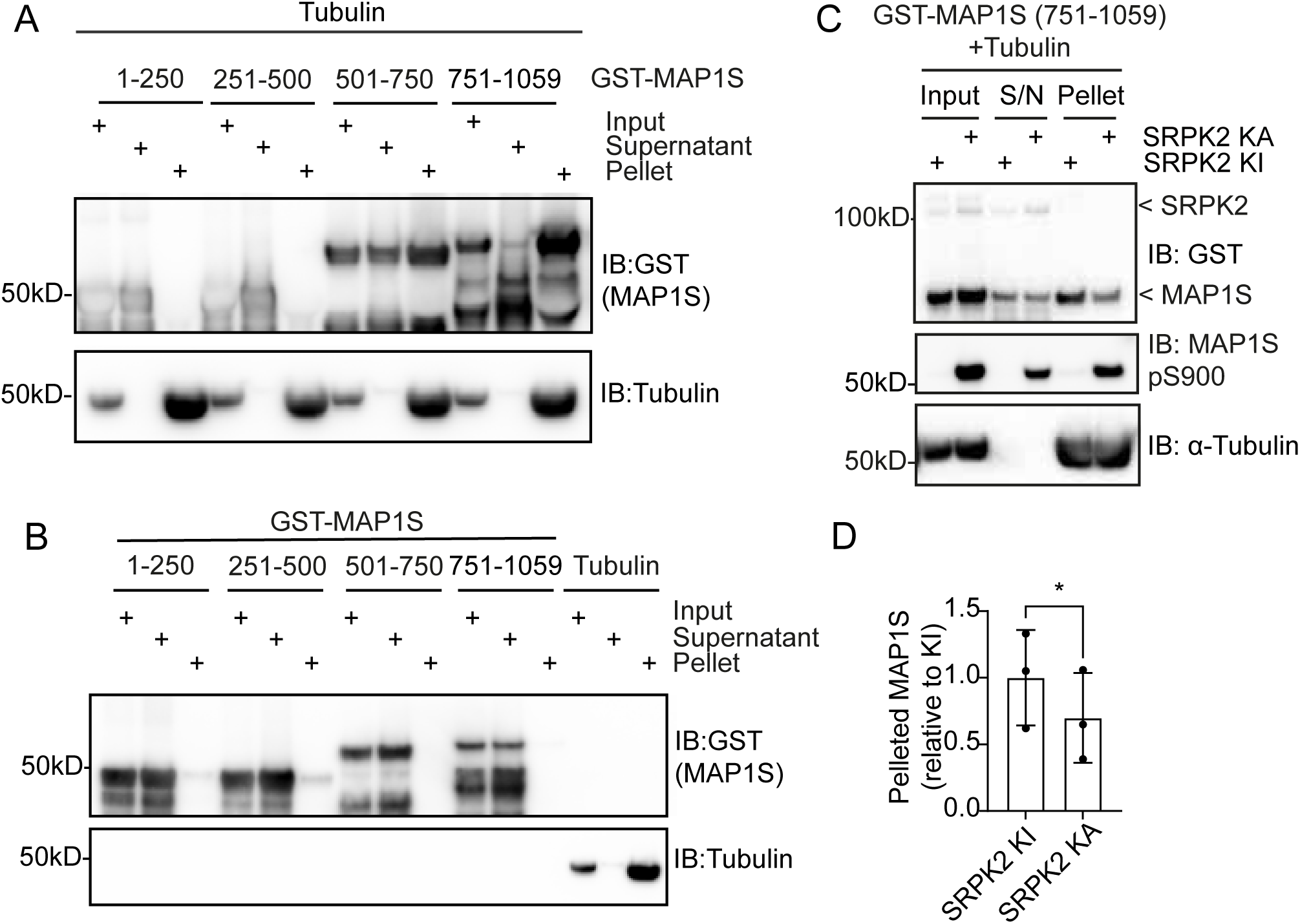
MAP1S-microtubule association is negatively regulated by SRPK (A) Co-sedimentation of 4 GST-hMAP1S fragments with microtubules after ultracentrifugation was analysed by immunoblotting (IB) with anti-GST and anti-tubulin antibodies (n=3). (B) Control experiment showing sedimentation of GST-hMAP1S fragments and polymerised tubulin (microtubules) after ultracentrifugation. Input, supernatant, and pellet fractions were analysed by immunoblotting (IB) for anti-GST and anti-tubulin (n=3). (C) Co-sedimentation of phosphorylated (kinase active SRPK, SRPK KA) and unphosphorylated (kinase inactive SRPK, SRPK KI) GST-hMAP1S (751-1059) with microtubules was analysed by immunoblotting (IB) for anti-GST, anti-MAP1S pS900, and anti-Tubulin antibodies (n=3). (D) Quantitative analysis of co-sedimentation of phosphorylated (SRPK KA) and unphosphorylated (SRPK KI) GST-hMAP1S (751-1059) with microtubules. The ratio of GST-hMAP1S (751-1059) that co-sediments with microtubules in the pellet to total GST-hMAP1S (751-1059) was determined. Data are presented as mean ± standard deviation (SD) (n=3) * = P < 0.05.

We next sought to investigate the impact of SRPK phosphorylation on MAP1S microtubule interaction. Incubation of GST-MAP1S aa751-1059 with active SRPK2 (KA), but not a kinase-inactive mutant (KI), leads to GST-MAP1S aa751-1059 phosphorylation, as measured by immunoblotting with an anti-MAP1S pS900 antibody (Fig. 4C). This prompted us to test whether SRPK phosphorylation affects the interaction of GST-MAP1S fragments with microtubules by co-sedimentation. Unphosphorylated GST-MAP1S aa751-1059 incubated with SRPK KI co-sediments with microtubules upon ultracentrifugation (Fig. 4C). However, phosphorylation by active SRPK WT inhibits co-sedimentation of GST-MAP1S aa751-1059 with assembled microtubules (Fig. 4C, D). Therefore, SRPK phosphorylation proximal to a major MAP1S microtubule binding region inhibits microtubule binding. This is consistent with previous reports that other kinases, such as CDKL5, can phosphorylate MAP1S and modulate microtubule binding in a similar manner ^47^.

### SRPK phosphorylation modulates proteolytic processing of MAP1S by CAPN10

Our finding that SRPK phosphorylates MAP1S at sites that flank the CAPN10 proteolytic cleavage site prompted us to address whether SRPK phosphorylation of MAP1S modulates proteolytic cleavage into HC and LC polypeptides. Thus, we generated a MAP1S mutant that cannot be phosphorylated by SRPK. Mouse MAP1S is phosphorylated in an SRPK-activity-dependent manner on 4 distinct motifs containing up to 10 phosphorylation sites (Fig. 2). Therefore, we substituted each of these Ser/Thr sites to non-phosphorylatable Ala (MAP1S S/T10A). We first tested whether MAP1S S/T10A disrupts MAP1S phosphorylation to a similar extent as SRPK inhibition. As shown previously, MAP1S WT runs as 2 species when analysed by Phos-tag immunoblot, whilst SRPK inhibition leads to the appearance of a species with increased mobility (Fig. 5A). MAP1S S/T10A also exhibits increased mobility on Phos-tag SDS-PAGE (Fig. 5A), suggesting a reduction in phosphorylation compared to MAP1S WT. Importantly, MAP1S S/T10A exhibits similar mobility to the least phosphorylated species observed upon SRPK inhibition in mESCs (Fig. 5A). However, a pool of MAP1S WT remains phosphorylated upon SRPK inhibition (Fig. 5A), suggesting that this pool may be protected from the action of phosphatases in cells. Nevertheless, our data indicate that MAP1S S/T10A abolishes SRPK-dependent phosphorylation of MAP1S.

**Figure 5:**
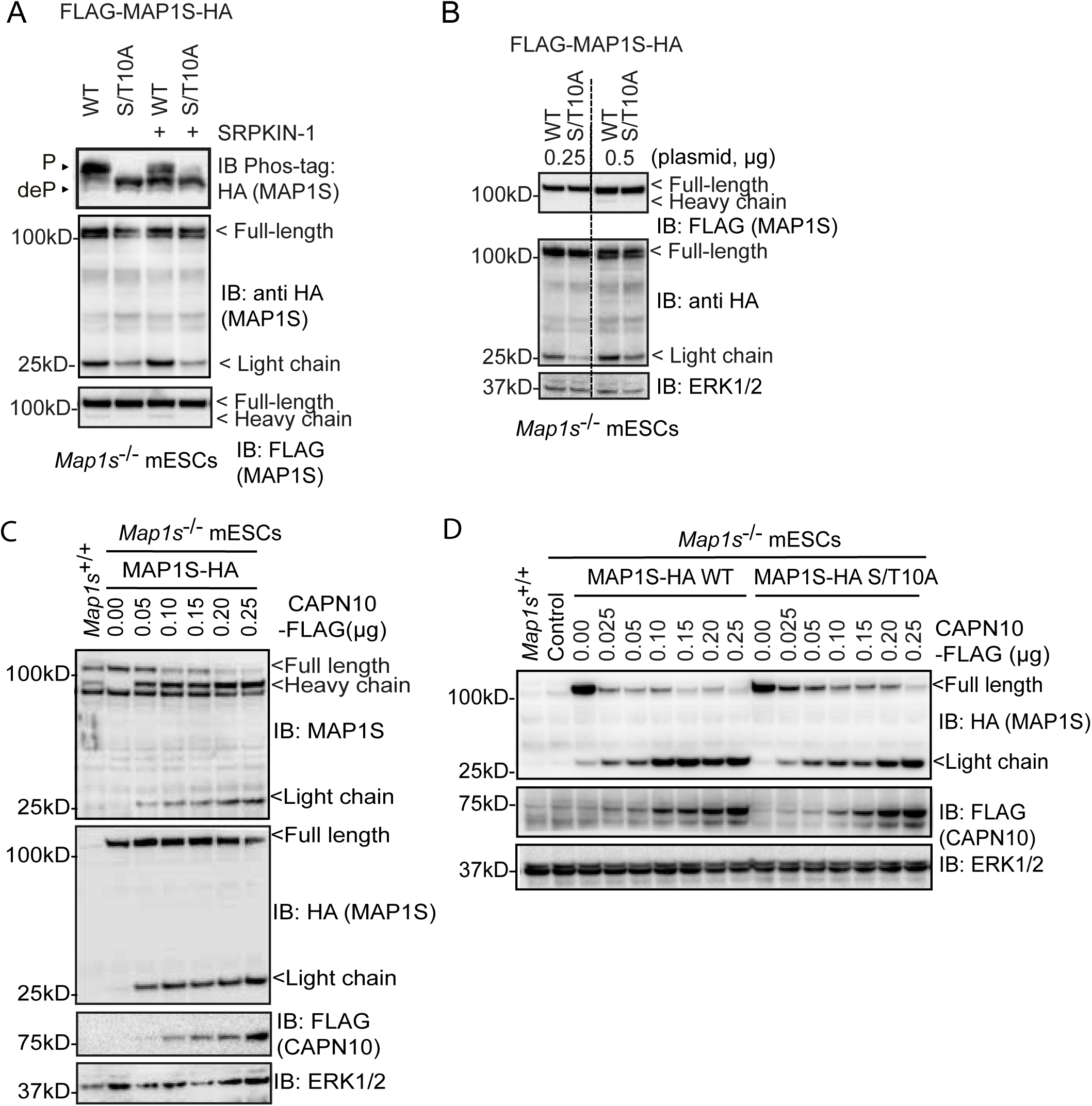
SRPK phosphorylation promotes CAPN10-mediated proteolysis (A) *Map1s*^-/-^ mESCs were transfected with FLAG-mMAP1S-HA or FLAG-mMAP1S-HA S/T10A, treated with DMSO or 2 µM SRPKIN-1 for 6 hours, and mMAP1S phosphorylation and expression analysed by Phos-tag immunoblotting for anti-HA and regular immunoblotting (IB) for anti-HA and anti-FLAG. P = phosphorylated MAP1S; deP = dephosphorylated MAP1S (n≥3). (B) *Map1s*^-/-^ mESCs were transfected with FLAG-mMAP1S-HA WT or S/T10A and analysed by immunoblotting (IB) for anti-FLAG and anti-HA. ERK1/2 was used as a loading control (n=3). (C) *Map1s*^-/-^ mESCs were co-transfected with mMAP1S-HA and increasing amounts of mouse Calpain-10-FLAG and analysed by immunoblotting (IB) for anti-MAP1S, anti-HA, and anti-FLAG. ERK1/2 was used as a loading control (n≥3). (D) *Map1s*^-/-^ mESCs were co-transfected with either mMAP1S-HA WT or S/T10A and increasing amounts of mouse CAPN10-Flag, and analysed by immunoblotting (IB) for anti-HA and anti-FLAG. ERK1/2 was used as a loading control (n=3). *Map1s*^+/+^ mESC lysate was included as a positive control, and empty vector (EV) as a negative control.

To test our hypothesis that SRPK phosphorylation of MAP1S impacts CAPN10-mediated proteolysis, we expressed WT and non-phosphorylatable S/T10A MAP1S and examined the processing of full-length (FL) MAP1S into HC and LC fragments. A dual N-and C-terminally tagged FLAG-MAP1S-HA construct was employed, where the N-terminal FLAG-tag can be used to visualise MAP1S FL and HC, and the C-terminal HA-tag can be used to visualise MAP1S FL and LC. In mESCs, MAP1S is relatively poorly processed into HC and LC fragments, as evidenced by the relatively low HC and LC signal in comparison to the FL signal in each case (Fig. 5A, B). However, MAP1S S/T10A exhibits reduced cleavage into HC and LC fragments (Fig. 5A, B), suggesting that SRPK phosphorylation of MAP1S may modulate CAPN10-mediated MAP1S cleavage.

As MAP1S is poorly processed into HC and LC fragments in mESCs, we sought to improve the processing by expressing CAPN10, which specifically catalyses proteolysis of the MAP1 family ^58^. As shown previously, in mESCs, MAP1S FL predominates, with low abundance of HC and LC fragments (Fig. 5B). However, increasing CAPN10 expression leads to decreased MAP1S FL abundance, and an increase in MAP1S HC and LC (Fig. 5C). Therefore, MAP1S proteolytic processing in mESCs can be driven by ectopic expression of CAPN10.

We then investigated whether MAP1S processing by overexpressed CAPN10 is modulated by SRPK-mediated MAP1S phosphorylation. Again, MAP1S FL is poorly processed in mESCs (Fig. 5B), and this is increased by CAPN10 expression (Fig. 5C). However, MAP1S S/T10A is poorly processed relative to MAP1S WT in both the presence and absence of overexpressed CAPN10 (Fig. 5D). These data suggest that SRPK phosphorylation of MAP1S at motifs surrounding the CAPN10 proteolytic cleavage site modulates processing of MAP1S FL into HC and LC fragments by CAPN10.

### MAP1S is progressively proteolysed during human neural differentiation

Proteolysis is thought to be required for the maturation of MAP1 family members into functional microtubule-binding proteins^58^, suggesting that this process may be important during neural differentiation. As MAP1S is poorly processed in pluripotent stem cells (mESCs; Fig. 5A, B), we hypothesised that MAP1S proteolysis is induced during neural differentiation, to enable formation of an active MAP1S microtubule binding complex that functions in e.g. axon formation in developing neurons. To test this, we exploited human induced pluripotent stem cells (hiPSCs) induced to differentiate to neuroepithelial cells (Day 6), neural progenitors (Day 12), and cortical neurons (Day 25; Fig. 6A). This is accompanied by suppression of pluripotency markers NANOG and OCT4, induction of neuroepithelial/neural progenitor markers SOX2 and PAX6, followed by neuronal markers TUJ1, TBR1, and FOXG1 (Fig. 6B). As expected, MAP1S is not efficiently cleaved into HC and LC in hiPSCs (Fig. 6C). However, as neural differentiation progresses, both MAP1S protein abundance and proteolytic processing increases such that most MAP1S FL is cleaved into HC and LC in Day 25 hiPSC-derived cortical neurons, (Fig. 6C). Our results therefore show that MAP1S proteolytic processing, which is modulated by SRPK phosphorylation and CAPN10, is induced upon differentiation of pluripotent stem cells to neurons.

**Figure 6:**
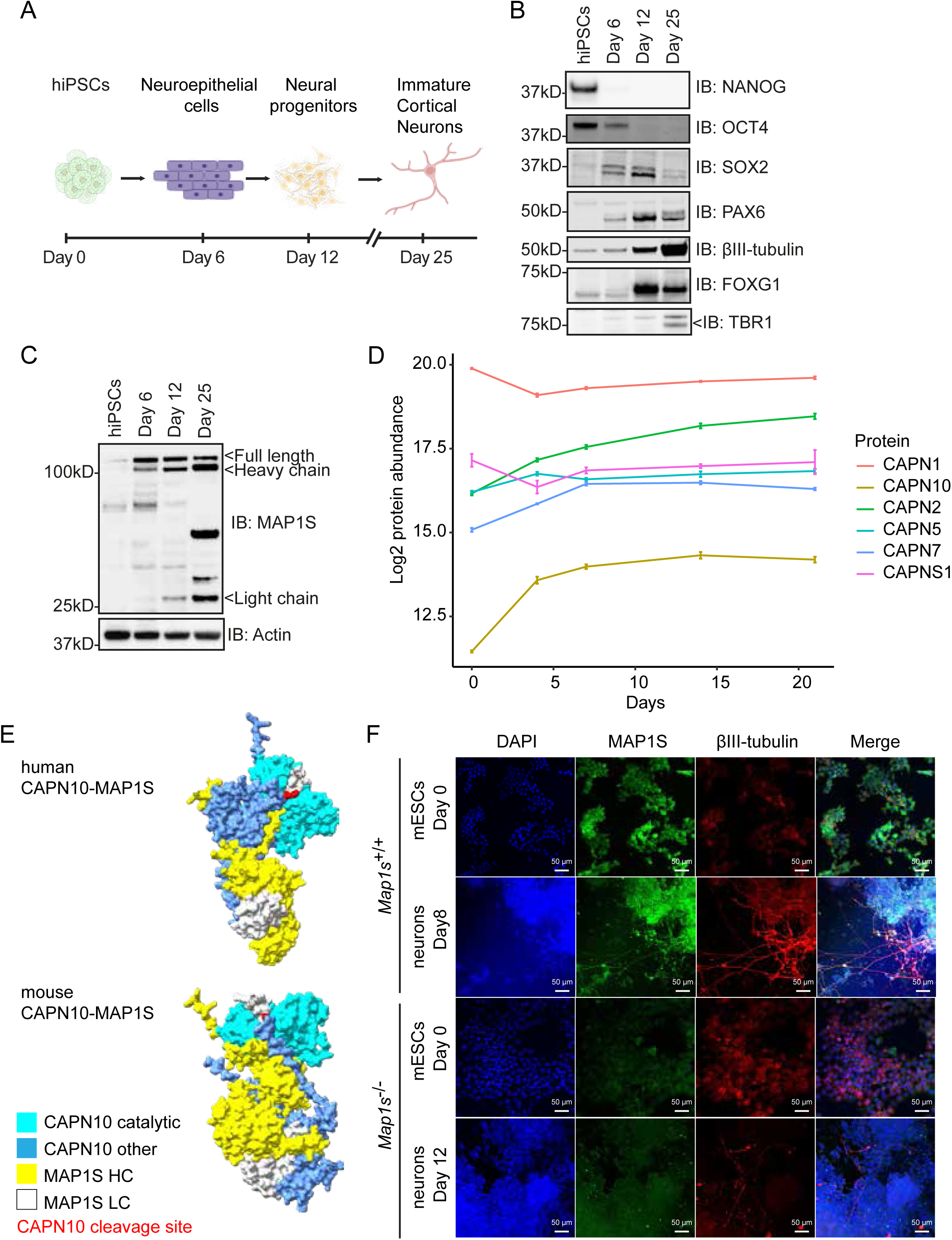
Progressive CAPN10 induction and MAP1S proteolysis during human neuronal differentiation (A) Schematic of stages during hiPSC cortical neuron differentiation. The figure was generated using BioRender. (B) hiPSCs were differentiated into cortical neurons and samples collected at day 0 (hiPSCs), day 6 (neuroepithelial cells), day 12 (neural progenitor cells), and day 25 (cortical neurons). The following markers were analysed by immunoblotting (IB): SOX2 = pluripotency/neural stem cells; NANOG and OCT4 = pluripotency; TBR1 = postmitotic cortical neurons; PAX6 = neural progenitors; βIII-tubulin = neurons; FOXG1 = early forebrain development (n=3). (C) hiPSCs were differentiated into cortical neurons, and samples were collected at day 0, day 6, day 12, and day 25, and analysed by immunoblotting (IB) for anti-MAP1S. Actin was used as a loading control (n=3). (D) Quantitative proteomic analysis of CAPN protein abundance during hiPSC (KOLF2.1J line) neural differentiation. Protein abundance of detected CAPN family members (CAPN1, CAPN2, CAPN5, CAPN7, CAPN10 and CAPNS1) was quantified at days 0, 4, 7, 14, 21, and 28 of neural differentiation. CAPN6 was inconsistently detected and quantified during neural differentiation and was excluded from the analysis. Graphs were generated by a visualization tool (https://niacard.shinyapps.io/Phosphoproteome/) using data from ^69^ (E) AlphaFold3 predictions of human and mouse MAP1S-CAPN10 complex (https://alphafoldserver.com/) analysed on ChimeraX 1.9 software. Predicted MAP1S disordered regions are not shown. Light blue = CAPN10 catalytic domain, blue = other regions of CAPN10, yellow = MAP1S heavy chain (HC), white = MAP1S light chain (LC), red = CAPN10 proteolytic cleavage site in MAP1S. (F) Mouse MAP1S localization in mESCs and mESC-derived neural cells was visualised by immunofluorescence microscopy of βIII-tubulin (red) and MAP1S (green). Nuclei were visualised by DAPI (blue). Scale bar = 50 μm (n ≥ 3).

### Dynamic expression of Calpain family proteases during human neural differentiation

To provide further insight into mechanisms by which MAP1S is regulated by proteolysis during human neural development, we used quantitative total proteomics analysis to assess the dynamics of CAPN abundance across a time course of hiPSCs differentiating to neurons ^69^. A total of 6 CAPN family members are consistently detected and protein abundances quantified during hiPSC neuronal development (Fig. 6D). Of those, CAPN1, CAPN5 and CAPNS1 remain largely constant over a time-course of human neural differentiation, while CAPN2 and CAPN7 increase slightly (Fig. 6D). However, CAPN10, which specifically cleaves MAP1S, dramatically increases in abundance during human neural differentiation (Fig. 6D), which correlates with processive MAP1S proteolysis observed during human cortical neuronal differentiation (Fig. 6C). Therefore, our data suggest that a CAPN10 expression switch acts in concert with SRPK phosphorylation to ensure efficient MAP1S proteolytic processing during human neurodevelopment.

### AlphaFold3 modelling provides a molecular basis for specific MAP1S proteolysis by CAPN10

Previous work suggests that MAP1S proteolysis into HC and LC fragments is specifically catalysed by CAPN10 ^58^. However, the molecular basis for this specificity has not been determined, and further CAPN family proteases are expressed at high levels during human neural differentiation (Fig. 6D). Therefore, we performed an *in silico* prediction of CAPN specificity for MAP1S cleavage into HC/LC using AlphaFold3. This was based on whether MAP1S i) is predicted to interact with CAPN family proteases that are expressed during human neurodevelopment, and ii) is predicted to interact in a configuration that supports MAP1S proteolytic cleavage at the CAPN cleavage motif. As proof-of-principle, our modelling indicates that both human and mouse CAPN10 interact with MAP1S in a manner that enables MAP1S proteolysis (Fig. 6E). However, while other CAPNs are predicted to interact with MAP1S (Fig. S2), and human CAPN1 and mouse CAPN2/5 interact with MAP1S in a manner predicted to enable proteolysis at the CAPN10 site (Fig. S2), only CAPN10 is predicted to complex with MAP1S in a manner that supports cleavage in both species (Fig. S2). In summary, our data indicate that CAPN10 expression is progressively induced during human neural differentiation, which likely drives specific cleavage of MAP1S into HC and LC fragments.

### MAP1S proteolysis during neural differentiation promotes microtubule interaction

Finally, we explored the physiological importance of MAP1S proteolytic processing to HC and LC. Previous studies have suggested that MAP1 family microtubule binding requires cleavage to HC and LC, either to enable assembly of these products into a MAP1 microtubule-binding complex ^58^, or for the cleaved MAP1 LC to interact with and stabilise microtubules ^56^. Therefore, we sought to investigate the co-localisation of MAP1S with microtubules in pluripotent cells and pluripotent stem cell-derived neurons. To this end, we took advantage of mESCs, to enable comparison of *Map1s*^+/+^ and *Map1s*^-/-^ cell lines and ensure the specificity of immunofluorescence staining. In mESCs, MAP1S localisation is mostly diffuse cytoplasmic, with little evidence of co-localisation with microtubules (Fig. 6F). Specificity of MAP1S staining is confirmed by *Map1s*^-/-^ mESCs (Fig. 6F). However, following MAP1S proteolytic processing during neural differentiation, MAP1S staining is colocalised with βIII-tubulin/TUJ1, a core microtubule component in neurons (Fig. 6F). These data therefore suggest that CAPN10-mediated processing of MAP1S into HC and LC fragments, which occurs during neural differentiation, may be important to enable MAP1S microtubule binding in neuronal cells.

## DISCUSSION

Neurological diseases are frequently characterised by dysregulation of the microtubule cytoskeleton, which is critical for neuronal integrity and functioning.

However, the precise mechanisms by which microtubule dynamics are regulated during the development of the nervous system remain poorly understood. In this study, we uncover a new layer of microtubule regulation by microtubule binding proteins, in which we identify Microtubule Associated Protein (MAP)1S as a novel substrate of Ser-Arg Protein Kinase (SRPK), which is itself implicated in neurological disorders. SRPK directly phosphorylates MAP1S at multiple sites in a C-terminal region involved in proteolytic maturation and microtubule binding, which modulates proteolytic processing of MAP1S by the CAPN10 protease and the affinity of the MAP1S microtubule binding domain for microtubules. MAP1S proteolytic processing is progressively induced during neurodevelopment, which appears to correspond with MAP1S acquisition of microtubule binding activity. Our results therefore demonstrate a key role for SRPK in enabling and controlling the maturation and function of a key microtubule regulatory protein during neurodevelopment, providing insight into mechanisms by which the microtubule cytoskeleton is regulated and potentially dysregulated in neurological conditions.

Future work will focus on establishing the mechanism by which MAP1S phosphorylation promotes CAPN10-mediated cleavage. Although AlphaFold modelling does not suggest direct contacts between SRPK phosphorylation sites on MAP1S and CAPN10, phosphorylation may nevertheless drive recruitment of MAP1S to CAPN10 to enable cleavage. Another possibility is that MAP1S phosphorylation modulates its sub-cellular localisation in proximity to CAPN10. Structural, biochemical and cell biology studies will unravel the role of MAP1S phosphorylation in driving CAPN10 recruitment and/or cleavage.

A key question concerns the role and regulation of MAP1S phosphorylation and proteolytic processing during neural development. In terms of regulation, we show that MAP1S is progressively proteolysed during neural specification. Our data also suggests that a pool of MAP1S is constitutively phosphorylated by SRPK in pluripotent stem cells, which in turn proposes that regulation of CAPN10 may be responsible for switching on MAP1S proteolysis. Indeed, quantitative proteomic analysis of hiPSCs during neural differentiation ^69^ indicates that CAPN10 dramatically increases in abundance. Therefore, we hypothesise that SRPK phosphorylation is a permissive requirement for efficient MAP1S proteolysis, which is then executed by CAPN10 induction during neuronal differentiation.

The functional role of MAP1S processing during neural differentiation also requires further study. Previous data suggest that proteolysis of MAP1 family proteins activates microtubule binding, either by releasing the light chain or by enabling complex formation between cleaved heavy and light chains ^58^. This may have implications for the development of neurons, particularly relating to processes that are dependent on microtubule dynamics, such as axon elongation, stabilisation, and dendritic spine formation.

Finally, future studies should focus on the importance of MAP1S phosphorylation and proteolytic processing in human disease. MAPs play a key role in neurological disorders, including neurodegeneration and intellectual disability ^44,45,49,50,70–73^, and aberrant phosphorylation has been implicated in dysregulation of microtubule binding activity and microtubule dynamics ^41,42,45,47^. Our discovery that MAP1S phosphorylation modulates proteolytic processing during neural development suggests that this mechanism may be disrupted in neurological disorders. Indeed, SRPK mutation/overexpression/deletion has been implicated in neurodegeneration, specifically Alzheimer’s disease ^44,74,75^, and intellectual disability/autism ^17–21^.

Therefore, dysregulation of MAP1S function could be at least partially responsible for etiology or neurological diseases via microtubule cytoskeleton dysfunction.

## AUTHOR CONTRIBUTIONS

ZY performed and analysed most of the experiments presented in Figs. 2, 3, 4, 5, and 6. FBustos and HZ performed the phosphoproteomic workflow and MG performed data analysis in Fig. 1. EKJH developed and validated the SRPK inhibitor-resistant mutant and provided Figure 1I. ML performed hiPSC neural differentiation and provided Figure 6A. IS assisted with data analysis. IW investigated *in vitro* phosphorylation of MAP1S fragments and performed 1 replicate of *in vitro* targeted phosphoproteomics. YX, HY, and YAQ performed total quantitative mass spectrometry and developed the data visualisation tool used in Fig. 6D. RG performed targeted mass spectrometry and assisted with data analysis. FBrown developed and purified the MAP1S phospho-S786 antibody. CJH helped with protein production. RT performed DNA cloning. TM performed CRISPR design and construct preparation. ZY and GMF assembled figures and wrote the paper.

## ACKNOWLEDGEMENTS

The authors would like to acknowledge Prof. John Rouse (University of Dundee) and Mark Dorward, Clare Johnson, Victoria Barlow, and Ffion Thomas (MRC-PPU Reagents & Services, University of Dundee) for purified GST-MAP1S fragments and generating anti-MAP1S-pS900 antibodies, and Dr. Renata Soares (MRC-PPU Mass Spectrometry, University of Dundee) for data handling and uploading raw mass spectrometry data to the EBI-PRIDE server. ZY was funded by a China Scholarship Council 4-year PhD studentship. ML, EKJH, and GMF were funded by a Wellcome Trust Discovery Award (225930/Z/22/Z). GMF was also funded by a Wellcome Trust/Royal Society Sir Henry Dale Fellowship (211209/Z/18/Z). This research was supported in part by the Intramural Research Program of the NIH, National Institute on Aging (NIA), National Institutes of Health, Department of Health and Human Services; project number ZIAAG000534. This research was supported by the Intramural Research Program of the National Institutes of Health (NIH). The contributions of the NIH author(s) are considered Works of the United States Government. The findings and conclusions presented in this paper are those of the author(s) and do not necessarily reflect the views of the NIH or the U.S. Department of Health and Human Services.

## DATA AVAILABILITY

Raw data from phosphoproteomic profiling mass spectrometry (Fig. 1) is deposited on EBI-PRIDE (https://www.ebi.ac.uk/pride), accession number PXD068224. Proteomic data from hiPSC neuronal differentiation is publicly available via the web tool https://niacard.shinyapps.io/Phosphoproteome/ ^69^. All other data is available by request.

## SUPPLEMENTAL INFORMATION

Document S1

Figures S1-S2 and associated figure legends

