## SUPPLEMENTAL MATERIALS for "Phosphorylation enables progressive microtubule-associated protein proteolysis and functionalisation during neural development"

**SUPPLEMENTARY INFORMATION**

**Figure S1**

Statistical analysis of Extracted Ion Chromatogram (XIC) values for abundance of MAP1S peptides following DMSO and SRPKIN-1 treatment. Phosphopeptides with the same sequence and predicted phosphorylation site(s) but different observed mass/charge ratios were analysed together. Data are presented as mean ± standard deviation (SD) for identified peptides of the indicated observed masses that contain the indicated phosphorylation sites. Statistical significance was determined by the Shapiro-Wilk test for normal distribution on GraphPad Prism 9.4.0. After normal distribution, the Wilcoxon Signed-Rank Test (for S684) or paired T-test (for S785/S786, S786, S458/S462, S462, S520, S684/687, S658/T659/S660, S716/S724, and S724) was used for comparing the difference between DMSO and SRPKIN-1 treatment groups on GraphPad Prism 9.4.0. ns= p>0.05, * = p<0.05, *** = p<0.001, **=P<0.01, **** = p<0.0001. MAP1S S462 and S687 were omitted from further functional analysis due to poor-quality XIC data from S687 phosphopeptides (shown in Fig. S1E), and because singly phosphorylated S462 phosphorylated peptides showed no significant difference between DMSO and SRPKIN-1.

**Figure S2**

AlphaFold3 predictions of human and mouse MAP1S-CAPN complexes (https://alphafoldserver.com/) were analysed on ChimeraX 1.9 software. Predicted MAP1S disordered regions are not shown. Light blue = CAPN10 catalytic domain, blue = other regions of CAPN10, yellow = MAP1S heavy chain (HC), white = MAP1S light chain (LC), red = MAP1S CAPN10 proteolytic cleavage site.

Figure S1

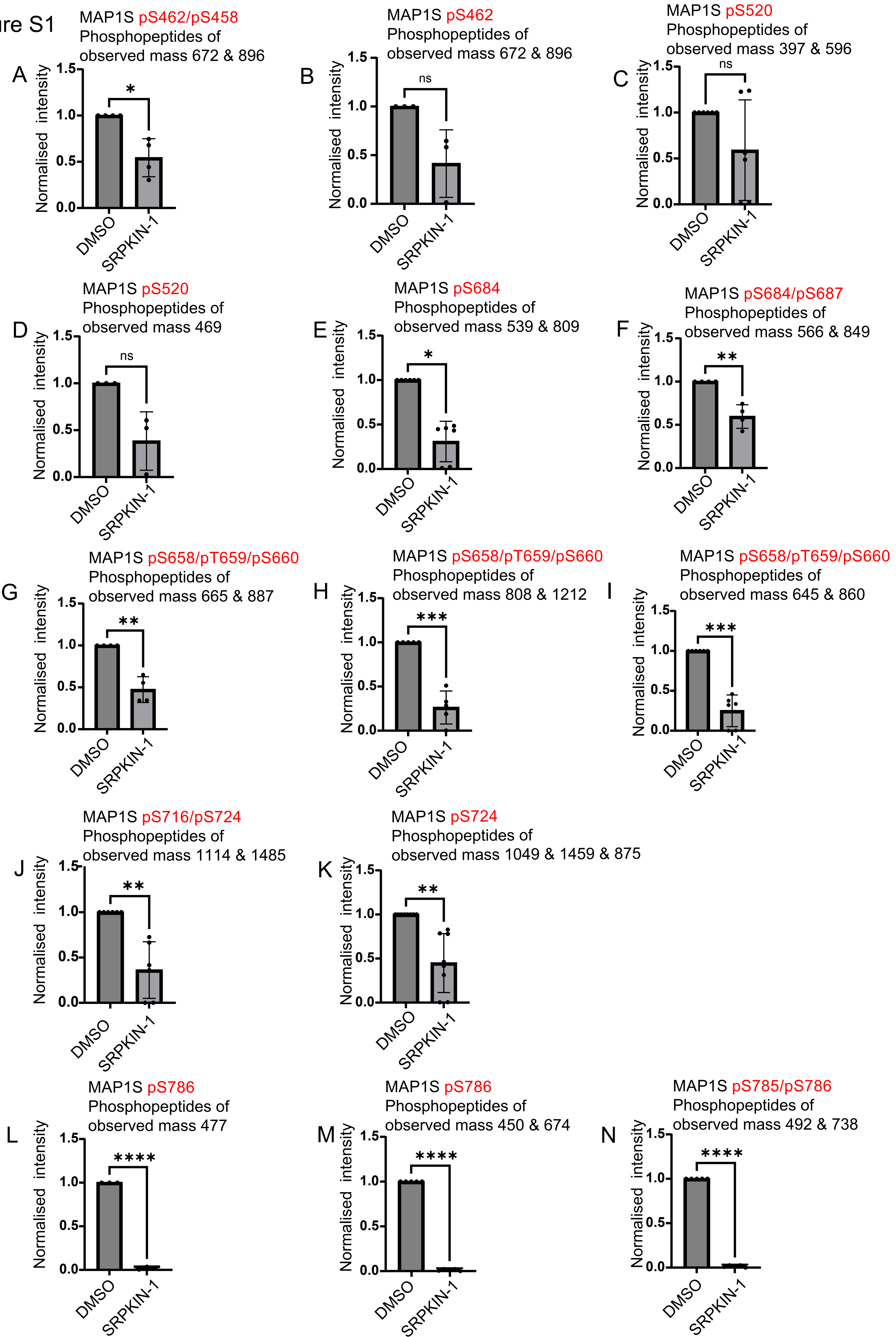

A
MAP1S pS462/pS458
Phosphopeptides of
observed mass 672 & 896
Normalised intensity
1.5
1.0
0.5
0.0
*
DMSO
SRPKIN-1
B
MAP1S pS462
Phosphopeptides of
observed mass 672 & 896
ns
C
MAP1S pS520
Phosphopeptides of
observed mass 397 & 596
ns
D
MAP1S pS520
Phosphopeptides of
observed mass 469
ns
E
MAP1S pS684
Phosphopeptides of
observed mass 539 & 809
*
F
MAP1S pS684/pS687
Phosphopeptides of
observed mass 566 & 849
**
G
MAP1S pS658/pT659/pS660
Phosphopeptides of
observed mass 665 & 887
**
H
MAP1S pS658/pT659/pS660
Phosphopeptides of
observed mass 808 & 1212
***
I
MAP1S pS658/pT659/pS660
Phosphopeptides of
observed mass 645 & 860
***
J
MAP1S pS716/pS724
Phosphopeptides of
observed mass 1114 & 1485
**
K
MAP1S pS724
Phosphopeptides of
observed mass 1049 & 1459 & 875
**
L
MAP1S pS786
Phosphopeptides of
observed mass 477
****
M
MAP1S pS786
Phosphopeptides of
observed mass 450 & 674
****
N
MAP1S pS785/pS786
Phosphopeptides of
observed mass 492 & 738
****

Figure S2

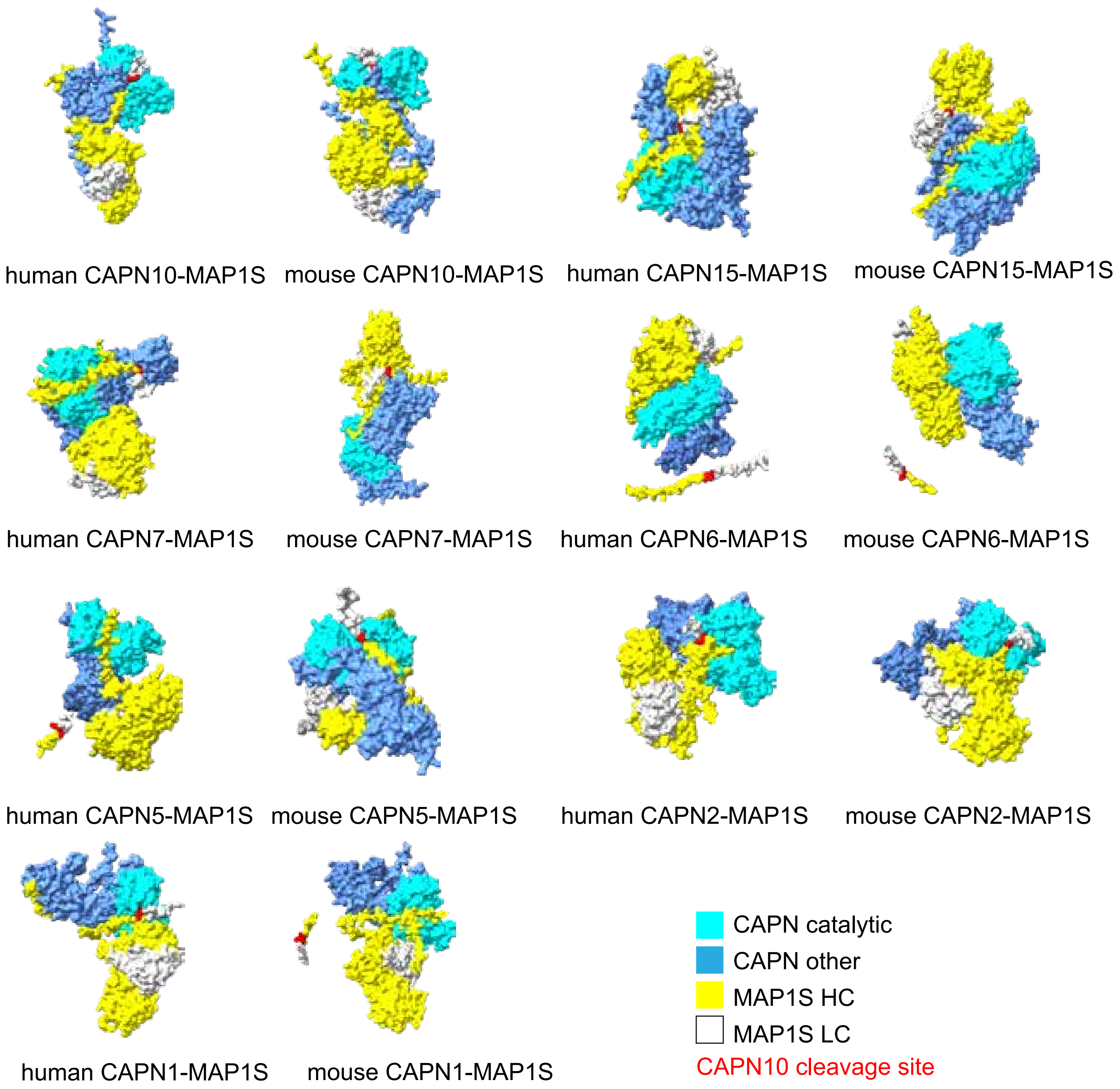
human CAPN10-MAP1S
mouse CAPN10-MAP1S
human CAPN15-MAP1S
mouse CAPN15-MAP1S
human CAPN7-MAP1S
mouse CAPN7-MAP1S
human CAPN6-MAP1S
mouse CAPN6-MAP1S
human CAPN5-MAP1S
mouse CAPN5-MAP1S
human CAPN2-MAP1S
mouse CAPN2-MAP1S
human CAPN1-MAP1S
mouse CAPN1-MAP1S
CAPN catalytic
CAPN other
MAP1S HC
MAP1S LC
CAPN10 cleavage site
